# A Conserved Mechanism for Positioning Ferredoxin–NADP⁺ Reductase at Photosystem I in Green Algae

**DOI:** 10.64898/2026.04.07.716946

**Authors:** Shira Artman, Pini Marco, Tamar Elman, Oren Ben-Zvi, Yoav Dan, Lihi Adler-Abramovich, Yuval Mazor, Iftach Yacoby

## Abstract

The association of ferredoxin-NADP^+^ reductase (FNR) with thylakoid membranes constitutes a central regulatory node in photosynthetic electron transport, governing NADPH production essential for carbon fixation. In cyanobacteria and higher plants, this interaction is mediated by an intrinsic FNR domain or specialized proteins, yet the mode of recruitment in green algae has remained enigmatic. Here, we show that in the green microalga *Chlamydomonas reinhardtii*, FNR is directly tethered to photosystem I-LHCI (PSI-LHCI) through a conserved N-terminal α-helix of the antenna protein Lhca4. Cryogenic electron microscopy localizes FNR to the stromal side of PSI proximal to Lhca4, while AlphaFold modeling identifies a specific interaction interface, which we validate using isothermal titration calorimetry. Structural modeling further reveals that the spatial separation between PSI-bound FNR and ferredoxin (Fd) is incompatible with direct electron transfer, indicating that it occurs sequentially rather than through a stable PSI-Fd-FNR complex. Comparative analysis demonstrates that the FNR-binding N-terminal motif of Lhca proteins is conserved across diverse green microalgae, suggesting an evolutionarily conserved strategy for positioning FNR at PSI. Collectively, our results uncover a novel mechanism for FNR recruitment and establish a new principle by which photosynthetic electron partitioning is regulated through spatial organization of electron transfer components.

## Introduction

Photosynthesis power life by producing energy-rich molecules and oxygen from sunlight thorough linear electron flow (LEF) in the thylakoid membrane, generating ATP and NADPH to sustain carbon fixation^1,2^. Electrons released from water splitting at photosystem II (PSII) are transferred via plastoquinone (PQ), cytochrome *b6f* (cyt *b6f*) and plastocyanin (PC) to photosystem I (PSI), where they are excited and delivered to ferredoxin (Fd)^3^. Electrons can then be diverted into several pathways, the preferred of which involves ferredoxin-NADP^+^ reductase (FNR), which reduces NADP^+^ to NADPH^4^. FNR is composed of a domain that binds its prosthetic group flavin adenine dinucleotide (FAD) and a NADP^+^-binding region^5,6^. LEF alone does not produce the ATP/NADPH ratio required for the Clavin-Benson-Bassham (CBB) cycle. This imbalance is compensated by modulating electron transport through cyclic electron flow (CEF) around PSI, which yields ATP without generating NADPH^7^. CEF can be carried out by the NADH dehydrogenase-like (NDH) complex or through a PGR5 and PGRL1-dependent route^8^.

PSI is composed of a core complex containing several protein subunits designated PsaA-PsaL, PsaN and PsaO^9^. In higher plants and green algae, PSI is associated with light harvesting complex I (LHCI), which transfers the absorbed light energy to the PSI core^10,11^. In the green microalgae *Chlamydomonas reinhardtii* (*C. reinhardtii*), LHCI comprises ten Lhca proteins that form three distinct belts: an inner belt (Lhca1_in_, 8, 7, and 3) associated with the PSI core, an outer belt (Lhca1_out_, 4, 6, and 5) surrounding the inner belt, and a side belt (Lhca2 and 9) also interacting with the PSI core^9,12,13^. The inner belt is equivalent to the LHCI of plants, which consists of only four Lhca proteins^14^. In contrast, cyanobacteria harvest light using phycobilisomes (PBSs), large non-membrane peripheral antenna complexes that primarily associate with PSII, facilitating the transfer of excitation energy to both PSII and PSI^15,16^. PBSs usually have an allophycocyanin-containing core, with radiating phycocyanin-rich rods, that are connected to each other by linker proteins, including cyanobacteria phycocyanin proteins G and D (CpcG and CpcD)^17^. In some cyanobacteria, there is an alternative PBS architecture in which a CpcG homolog termed CpcL enables it to bind to PSI^18^. One of the mechanisms that regulated the flow of electrons between cyclic and linear pathways is the association of FNR with thylakoids membranes^19^. This association occurs in distinct mechanisms in the different photosynthetic lineages. In higher plants, FNR binding is mediated by anchoring proteins, including Tic62 (translocon at the inner envelope of chloroplast) that binds FNR without obstructing its catalytic site and is thought to shield it from degradation in the dark^20,21^. TROL (thylakoid rhodanase-like protein) anchors FNR directly to the thylakoid membranes and is essential for maintaining efficient electron transport, particularly under high light intensities^21,22^. Both proteins contain a conserved FNR-binding region rich in Serine and Proline that confers a polyproline type II helix and binds FNR preferentially at low pH, enabling dynamic, pH-dependent regulation of FNR membrane association^23^. Another FNR-anchoring protein is LIR1, which forms complexes with FNR together with Tic62 and TROL. However, its absence had no impact on photosynthetic efficiency in *Arabidopsis thaliana* and only a modest effect in rice^24^. Together, these studies establish a plant-specific strategy in which FNR membrane localization is mainly governed by specialized proteins.

In cyanobacteria, the binding of FNR to thylakoid membranes relies mainly on a different mechanism involving a three-domain FNR containing an N-terminal CpcD-like domain in addition to the conserved FAD and NADP^+^ binding regions, which links FNR to PBSs^25^. Removal of this domain in *Synechocystis* sp. PCC 6803 terminated salt shock-induced CEF without impairing LEF, demonstrating that membrane binding of FNR is specifically required for CEF^26^. In addition, FNR lacking this N-terminal domain was shown to accumulate as free enzyme during iron or nitrate limiting conditions^27^. Association of this FNR isoform with CpcL-containing PBSs was shown to be essential for electron donation from NADPH to the PQ pool during photoheterotrophic growth^28^. Thus, cyanobacteria regulate FNR localization through structural features of FNR itself and antenna-associated interactions rather than dedicated proteins.

While FNR binding to thylakoid membranes has been extensively characterized in higher plants and cyanobacteria, the mechanisms operating in *C. reinhardtii* and other algae are not yet fully understood. In contrast to higher plants, *C. reinhardtii* lacks any known FNR-binding proteins^24,29^. In this organism, FNR association with thylakoids has been observed primarily under anaerobic conditions, where it is incorporated into super-complexes containing PSI-LHCI, LHCII, Cyt b6f, PGRL1, and additional factors, formed to support CEF and decrease LEF^30,31^. Structural analysis revealed that these super-complexes lack Lhca2 and Lhca9 side-belt subunits, and deletion of Lhca2 resulted in constant elevated CEF^31^. PGR5 and PGRL1 were shown to contributed to FNR association with PSI, as loss of either protein or both substantially reduced, yet did not abolish, PSI-bound FNR^29^, suggesting the presence of additional, PGR5/PGRL1-independent binding event. Supporting this notion, our previous work^32^ demonstrated direct interaction between FNR and PSI-LHCI that requires the presence of LHCI, yet not PGR5 or PGRL1. In addition, FNR did not exhibit binding to non-mature PSI, lacking LHCI, PsaF, PsaG and PsaK.

Here, we employed cryogenic electron microscopy (Cryo EM) of PSI-LHCI in complex with FNR and Fd to localize FNR to the vicinity of Lhca4. Inspection of Lhca4 sequences revealed a flexible N-terminal helix, absent from all PSI-LHCI published structures, yet verified by direct N-terminal sequencing of all mature antennae subunits in *C. reinhardtii* PSI-LHCI^33^. AF modeling suggested direct binding between this N-terminal Lhca4 domain and FNR. Isothermal titration calorimetry (ITC) confirmed the binding between Lhca4 N-terminus and FNR, which closely resembles the measured binding between purified PSI-LHCI and FNR^32^. Comparative sequence analysis revealed that this Lhca4 motif is conserved across green microalgae, yet absent from higher plants and cyanobacteria, establishing direct FNR binding to PSI-LHCI as an evolutionary distinct thylakoid membrane association with important implication for electron partitioning.

## Results

### Structural analysis indicates FNR binding at the N-terminus of Lhca4

We previously showed that in *C. reinhardtii*, FNR binds to the mature complex of PSI-LHCI at a 1:1 ratio, yet not to non-mature PSI devoid of its LHCI, PsaF, PsaG and PsaK, indicating that this binding takes place likely via these components^32^. To gain molecular insights of this interaction, we carried out Cryo EM analysis of *C. reinhardtii* PSI-LHCI incubated with both Fd and FNR at pH 7.5 and 6.5. In the resulting structures (Fig. 1A), PSI-LHCI was resolved to a global resolution of 2.2 and 2.29 Å at both pH conditions, showing a clear density for Fd (Fig. S1, S2 and Table S1). In addition, both maps showed a diffuse density just above Lhca4 at the stromal side of the complex which we attributed to FNR.

**Fig. 1.**
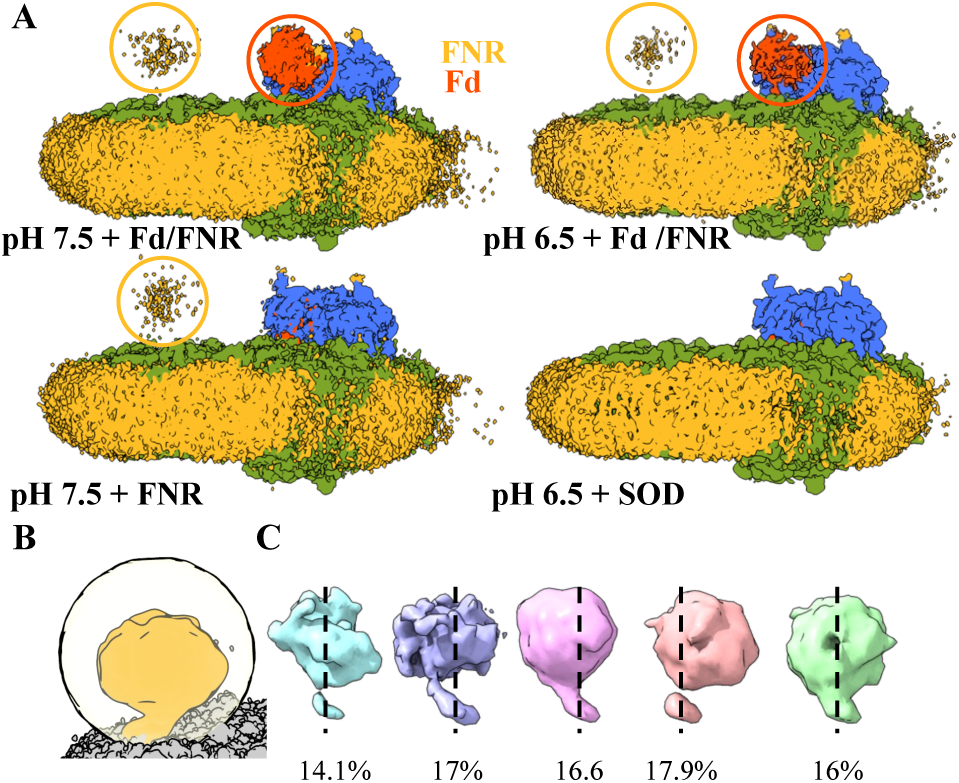
A PSI proximal, diffuse density, is associated with FNR in green Alga. (**A**) Cryo EM maps of PSI-LHCI from *C. reinhardtii* obtained from samples containing FNR and Fd at two different pH values (top) or containing FNR or SOD as the sole additive (bottom). (**B**) The FNR associated signal after subtracting the PSI-LHCI signal and renormalization (signal in gold, surrounded by the outline of the mask used in signal subtraction). (**C**) Results from 3D classification on the FNR associated signal showed classes oriented approximately evenly over the entire volume, consistence with a random distribution of FNR. The center of each frame is marked with a dotted line for reference.

To confirm that this density is associated with the presence of FNR, we carried out two control experiments. First, we imaged PSI-LHCI incubated with Superoxide dismutase (SOD) but without Fd and FNR. In the resulting 2.92 Å resolution Cryo EM map, we observed no additional densities in the Fd binding site or above Lhca4 (Fig. 1A and Fig. S3). In addition, we carried out experiments in which only FNR was incubated with PSI-LHCI. These maps were obtained at a resolution of 2.5 Å and clearly showed a diffuse density above Lhca4, confirming the association of this density with the presence of FNR in the reaction (Fig. 1A and Fig. S4).

Subtraction of the PSI signal and subsequent renormalization highlighted the FNR-associated signal (Fig. 1B). 3D classification of this signal indicated that it adopts multiple orientations (Fig. 1C, classes #1–#5). In all classes, a volume roughly the size of FNR, accompanied by a “tail” connecting it to the PSI stromal side, is seen. Each class consists of a similar fraction of particles (∼14–18%), suggesting a random orientation for FNR within this volume. Due to its small size of ∼35 kDa^5^, which is below the lower limit of detection for Cryo EM (∼38 kDa)^34^, it was not possible to reconstruct a high-resolution map showing FNR directly. Hence, we further analyzed the appearance of this density using AlphaFold (AF) to model the interaction between FNR and Lhca4.

Close examination of the Lhca4 gene from *C. reinhardtii* revealed a short sequence (24 amino acids) following the chloroplast import signal sequence and before the first amino acids observed in the Cryo EM map unique to Lhca4, hence this portion of Lhca4 is absent from both current and previously published structures^12,13,35,36^. The presence of this peptide in the mature protein was previously verified using N-terminal sequencing of all the mature Lhca proteins composing LHCI of *C. reinhardtii*^33^.

When the full sequence of Lhca4, including this N-terminal extension, was submitted as input for AF, a putative complex between the two proteins was generated (Fig. 2A). The model contains a novel interface formed by the N-terminal region of Lhca4, which folds as an α-helix and interacts with a cleft in FNR (Fig. 2A). The AF model immediately suggests an interpretation for the diffuse density observe above Lhca4 in the Cryo EM samples that included FNR. All the AF output models positioned the Lhca4 helix at the same FNR binding pocket, yet differed in the relative orientation between FNR and Lhca4, suggesting a flexible tether between the two proteins and consistent with the lack of distinct classes in the cryo EM data. The predicted structure of FNR closely conforms to the known FNR structure (Fig. 2A) comprising an N-terminal antiparallel β-barrel, followed by an extended loop and an α-helix. This N-terminal region binds the flavin adenine dinucleotide (FAD) cofactor, while the C-terminal region, consisting of a parallel β-sheet with several α-helixes arranged around it, binds NADP^+21,37–39.^ Similarly, the predicted structure of Lhca4 displays the conserved arrangement of Lhca proteins – three transmembrane helices, connected by shorter helical segments^13,40^. The AF model demonstrates high reliability, with a mean predicted local distance difference test (pLDDT) score of 86.85 indicating generally strong per-residue confidence^41^, and a predicted template modeling (pTM) score of 0.63, reflecting credible overall fold accuracy^42^. Notably, the N-terminal region of Lhca4 is modeled with a high mean pLDDT score of 84.19. The predicted aligned error (PAE) plot further supports high model confidence, showing low error values for most residues of both FNR and Lhca4, particularly at their interface (Fig. S8)^43^.

**Fig. 2.**
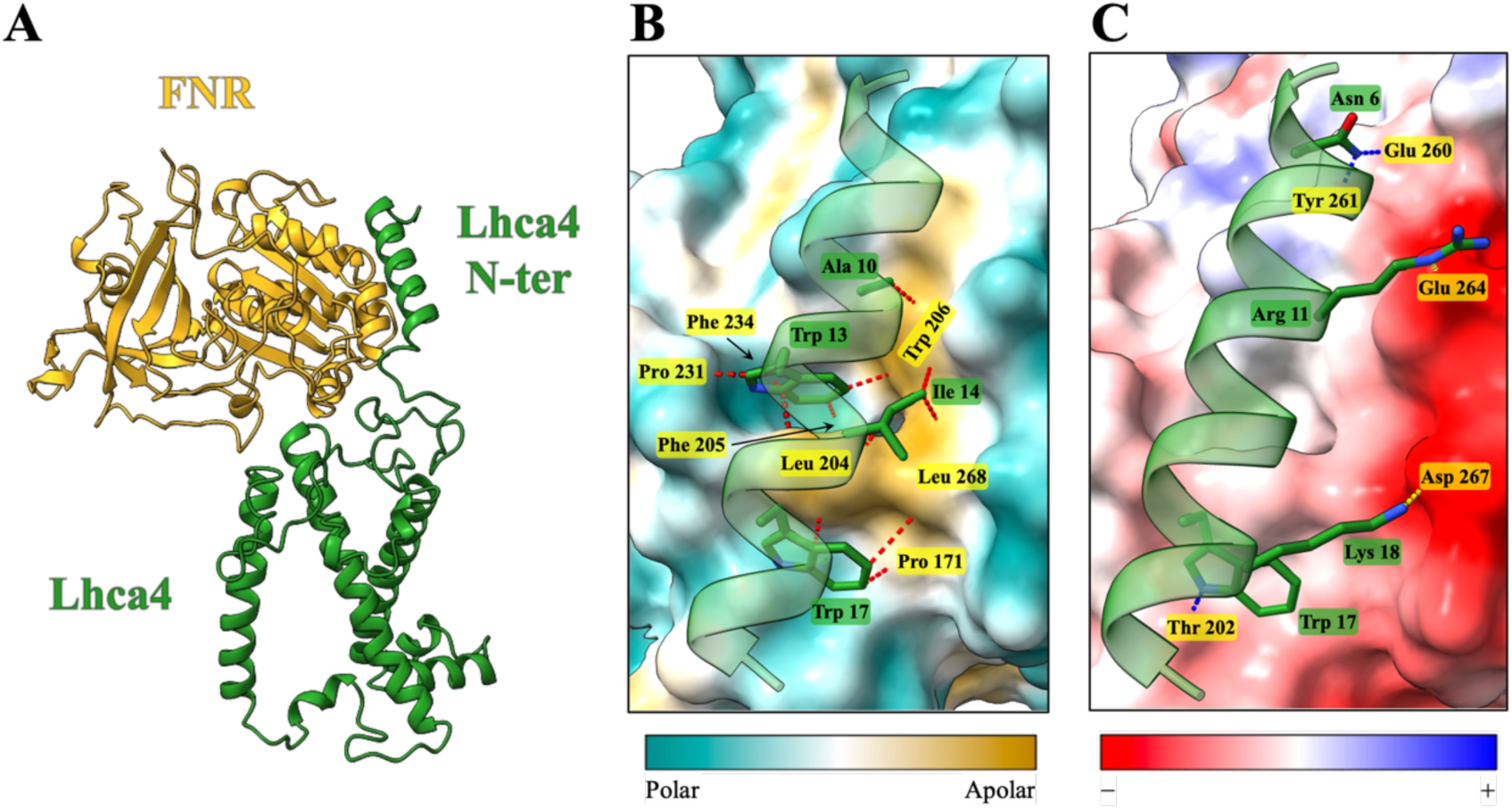
Modeling the interaction between FNR and Lhca4 in *C. reinhardtii*. **(A)** AF model of FNR (gold) and Lhca4 (dark green), showing the N-terminal helix of Lhca4 oriented toward FNR. **(B-C)** Detailed view of the interactions between the N-terminal helix of Lhca4 and FNR. Lhca4 residues are highlighted in green and FNR residues are highlighted in yellow. Only the shortest distances between each pair of residues are displayed. **(B)** Hydrophobic and van der Waals interactions (all distances <5 Å). FNR surface coloring indicates hydrophobicity, with cyan denoting hydrophilic (polar) regions and golden-brown denoting hydrophobic (apolar) regions. **(C)** Polar interactions (all distances <3 Å), with hydrogen bonds shown in blue and salt bridges in yellow. The surface of FNR is colored by electrostatic surface potential, ranging from red (negative) to blue (positive).

The interaction between FNR and Lhca4 is centered at the N-terminal helix of Lhca4 and is dominated by a hydrophobic patch and van der Waals contacts, with additional polar interactions, including hydrogen bonds and salt bridges outside this patch (Fig. 2B-C, Fig. S9). Apart from Trp 70, none of the interacting FNR residues are located within the FAD-binding domain or overlap with the NADP^+^ binding site. In accordance with these findings, sequence alignment of *C. reinhardtii* FNR together with FNR from *Anabaena* sp. PCC 7120 and *Zea mays* leaf and root isoforms indicates that these residues do not correspond to known Fd-binding sites (Fig. S10)^37,39,44,45^. The AF model also enabled estimation of the FNR-Lhca4 interaction interface, calculated either with the N-terminal helix or with the full Lhca4 protein, yielding buried surface areas of 611.1 Å^2^ and 1156.6 Å^2^, respectively. Based on the commonly applied estimate of ∼1 water molecule released per 10-12 Å^2^ of buried surface^46^, these interactions are expected to displace approximately 60 and 115 water molecules.

### Experimental confirmation of the predicted FNR-Lhca4 interaction site

To validate the putative interaction between FNR and Lhca4, a synthetic peptide corresponding to the N-terminal region of Lhca4 was synthesized and solubilized in aqueous buffer (Fig. 3A). The secondary structure of the solvated peptide was assessed using circular dichroism (CD) spectroscopy. The peptide exhibits a maxima peak at around 190 nm, and two minimum points at about 208 and 222 nm, typical to a CD profile of a α-helix (Fig. S11)^47^. Once it was verified that the peptide confers a α-helix, matching the AF-predicted structure of the N-terminus of the Lhca4 protein, its interaction with purified FNR was analyzed by isothermal titration calorimetry (ITC). As a control, a “charge-swapped” variant in which the acidic (negatively charged) residues were replaced with basic (positively charged) residues was synthesized and tested under identical conditions. This substitution is predicted to orient the N-terminal helix away from FNR (Fig. 3B), thereby disrupting their interaction. The mutated peptide isoform is also readily solubilized in aquas buffer and partially folds into α-helix (Fig. S11). The ITC results show that the native peptide exhibits strong, exothermic binding to FNR, as reflected by deep, negative injection peaks in the raw heat signal trace that cease at the end of the titration and a clear saturation plateau in the binding isotherm (Fig. 3C), enabling reliable determination of the thermodynamic parameters of the interaction, including the number of peptide molecules bound per FNR (N; 1.07 ± 0.03) and the dissociation constant (K_D_; 125.75 ± 58.17 nM). In contrast, the mutated peptide exhibits weaker binding to FNR – its raw heat signal trace peaks are noticeably shallower, persisting even at the end of the titration, thereby preventing the binding curve from reaching saturation at the same concentrations as the native peptide (Fig. 3D). Because saturation is not achieved, accurate thermodynamic parameters cannot be calculated for the mutated peptide. Nevertheless, these findings indicate that charge substitutions in the N-terminal region of Lhca4 substantially weaken its binding to FNR.

**Fig. 3.**
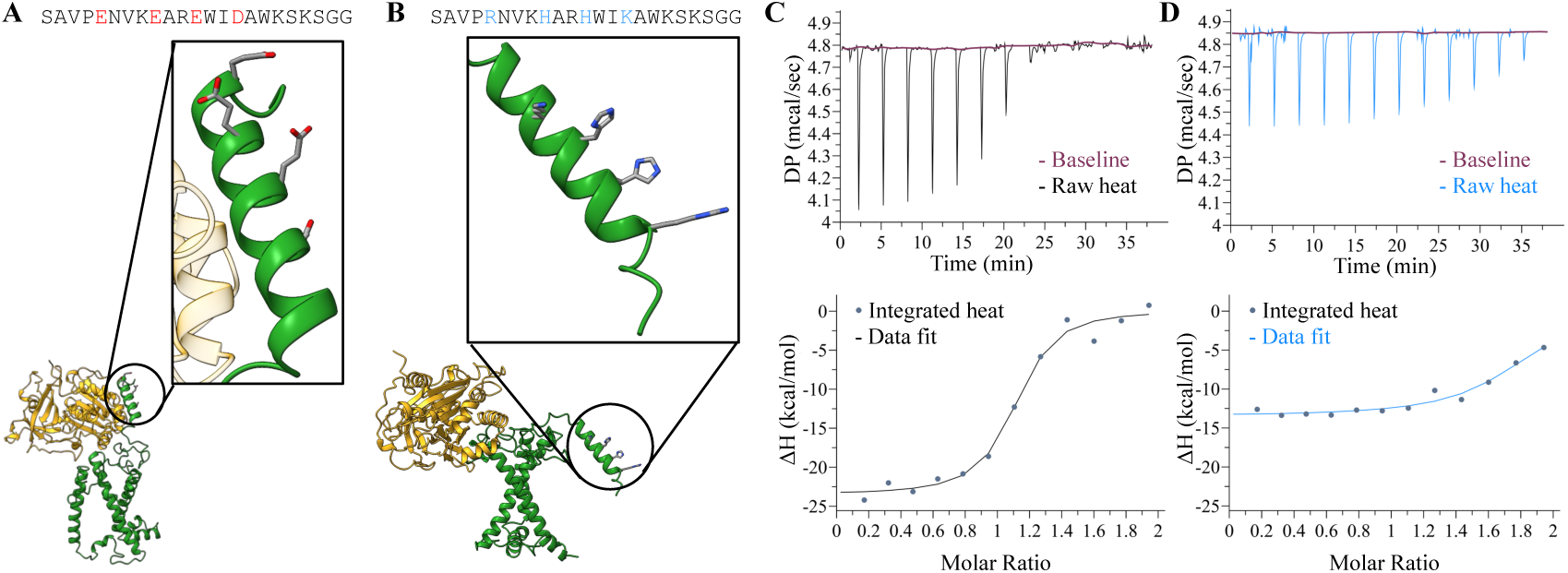
ITC analysis of FNR binding to native and mutant Lhca4-derived peptides. **(A-B)** Sequences and AF predicted structures of the native **(A)** and mutated **(B)** Lhca4-derived peptides, shown within the entire Lhca4 protein (green) and in complex with FNR (gold). Confidence parameters of the models are provided in Fig. S8 and S12, respectively. **(C-D)** Representative ITC experiments performed with the native **(C)** or mutated **(D)** peptides. Upper panels display raw heat signals recorded during titration of 100 μM peptide into a cell containing 10 μM FNR, with the purple line denoting the adjusted baseline. Lower panels present the corresponding integrated heat per injection (blue-grey points), with fitted curves obtained using a one-set-of-sites binding model implemented in MicroCal PEAQ-ITC Analysis Software. All experiments were conducted at 25°C in 50 mM Tris-HCl, pH 8. Averaging four independent measurements with the native peptide yielded an enthalpy change (ΔH) of −23.23 ± 0.97 kcal/mol, an entropy contribution (-TΔS) of 13.83 ± 0.46 kcal/mol and a free energy (ΔG) of −9.4 ± 0.11 kcal/mol.

### Fd to FNR electron transport is not likely when both are bound to PSI-LHCI

Next, we asked whether Fd could transfer electrons to FNR when both are bound to PSI-LHCI. To address this, we aligned the AF model of FNR and Lhca4 to our Fd-PSI-LHCI structure using Lhca4 as a reference, however this superimposition produced steric clashes. Therefore, we remodeled FNR with the outer LHCI belt (composed of Lhca5, 6, 4 and 1) in AF and repeated the alignment (Fig. S13). In the previously reported Fd-FNR complexes, the FAD group in FNR faces the Fd iron-sulfur cluster^48^, while in the AF model, it is oriented away from Fd.

The distance between the cofactors of Fd and FNR is 73.38 Å (Fig. 4), far exceeding the distance that is compatible with efficient electron transfer (10-20 Å)^49^. Notably, even the shortest atom-atom distance between Fd and FNR (23.13 Å), approximating the outer limit for the possible orientation of FNR, remains above this threshold (Fig. 4). Therefore, Fd is unlikely to reduce FNR while both proteins are simultaneously bound to PSI-LHCI. However, the binding site of the N-terminal helix of Lhca4 does not block the Fd binding site on FNR, making the PSI-bound FNR compatible with Fd binding, once Fd is released from PSI.

**Fig. 4.**
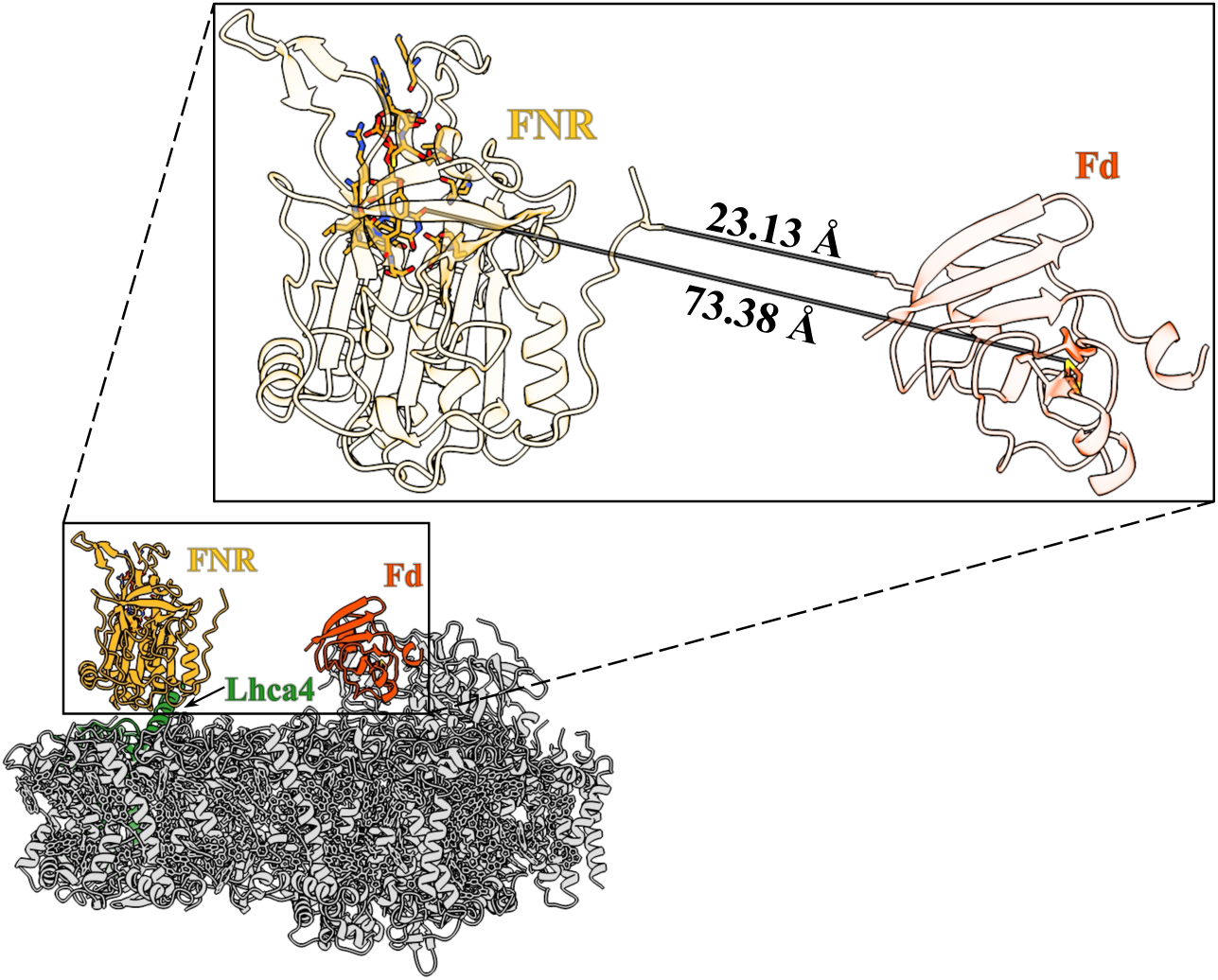
Spatial proximity of FNR and Fd associated with PSI-LHCI. The AF model of FNR (gold) in complex with the LHCI outer belt consisting of Lhca1, 4 (green), 6 and 5, was superimposed onto Fd (red) bound to PSI-LHCI (grey). To include the FAD cofactor, a root-type FNR structure (PDB 5H59) was superimposed onto FNR of the AF model. The minimal distance between FNR and Fd is 23.13 Å, whereas the closest distance between their cofactors is 73.38 Å.

### The FNR-binding helix is conserved across green microalgae

To evaluate the evolutionary conservation of the Lhca4-FNR interaction, we searched for Lhca proteins that contain an N-terminal extension across cyanobacteria and eukaryotes using Lhca4 from *C. reinhardtii* as query, with particular emphasis on those who aligned to its first 21 amino acids and were predicted to confer an α-helical conformation (Fig. 5A). FNR homologs were also detected across the identified taxa, generating species specific Lhca-FNR pairs as input for AF.

**Fig. 5.**
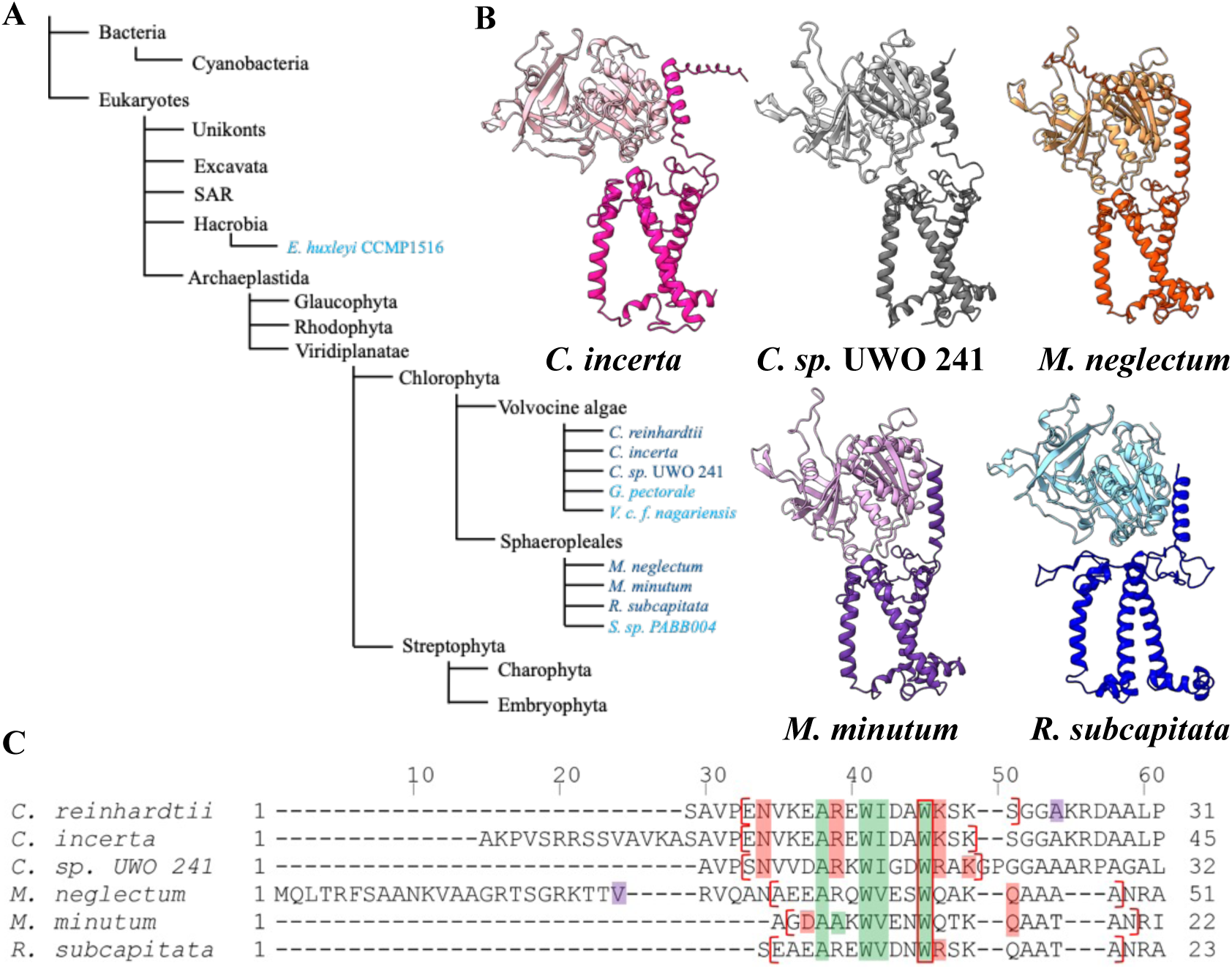
The interaction between FNR and the N-terminal helix of Lhca proteins is conserved in green microalgae. **(A)** Schematic phylogenetic tree (based on^53,54^) depicting the groups in which FNR-Lhca4 homologous interactions were searched. Dark blue indicates organisms with a high-confidence AF model, and light blue indicates those with a low-confidence model. **(B)** AF high-confidence models are shown. Each model includes the Lhca protein (darker shade) and its matching FNR sequence (lighter shade) retrieved from each species (*C. incerta*, pink; *C. sp.* UWO 241, grey; *M. neglectum*, orange; *M. minutum*, purple; *R. subcapitata*, blue). **(C)** Sequence alignment of the N-terminal regions of the Lhca proteins. Residues involved in hydrophobic interactions and van der Waals contacts are highlighted in green when located within the N-terminal helix and in purple when located outside it. Residues participating in polar interactions are highlighted in red. Residues involved in multiple types of interactions are framed according to their additional role. The residues forming the N-terminal helices are indicated by red brackets. The full Lhca proteins alignment is displayed in Fig. S22.

Models with sufficient confidence scores of pTM > 0.5 and ipTM > 0.8^42,50^ all show an interaction similar to the one observed between Lhca4 and FNR from *C. reinhardtii*, and exclusively include other green microalgae: *Chlamydomonas incerta*, *Chlamydomonas sp.* UWO 241, *Monoraphidium neglectum*, *Monoraphidium minutum* and *Raphidocelis subcapitata* (Fig. 5B). The modeled proteins match the typical structures of FNR and Lhca proteins, as also observed in the *C. reinhardtii* AF model described above. Every model also includes a protruding N-terminal helix, such as the one observed in Lhca4 of *C. reinhardtii*. To our knowledge, such a feature has not been described for Lhca proteins in any PSI-LHCI structures reported to date.

The PAE plots for *C. sp.* UWO 241, *M. minutum* and *R. subcapitata* support the validity of the interaction between the N-terminal helices of their Lhca subunits and their FNR proteins, as the area of interaction between these regions has low error values (Fig. S14). *C. incerta* and *M. neglectum*, however, seemingly possess an N-terminal extension that does not adopt an α-helical conformation (Fig. 5B), likely due to inaccurate signal peptide prediction by TargetP and SignalP, which failed to detect any signal peptide (Fig. S6). Their PAE plots (Fig. S14) reveal increased uncertainty in the relative positioning of these residues with respect to FNR, whereas the interaction involving the downstream N-terminal helix is associated with low PAE values, indicating a high-confidence predicted interface. The high reliability of these models is further supported by pTM and ipTM scores exceeding 0.5 and 0.8, respectively, together with mean pLDDT scores above 70, including for the N-terminal helices themselves (Fig. S15 and Table S4).

Organisms of other groups were also identified yet were excluded as they failed to meet the required criteria as described above (and in the Methods section). An exception is *Tetrabaena socialis*, a green microalga, for which only proteins with non-FNR annotations were identified, preventing prediction of interactions with its Lhca protein. It is noteworthy that *C. reinhardtii* Lhca4 homologs were not detected in either cyanobacteria or Unikonts (Fig. 5A). Other green microalgae met all the required criteria except their ipTM score was below 0.8 (Table S4), making them less reliable for further analyses. These include *Gonium pectoral*, *Volvox carteri f. nagariensis* and *Scenedesmus* sp. PABB004. Surprisingly, *Emiliania huxleyi* CCMP1516, a coccolithophore, also answered all criteria except for ipTM score (Table S4). Their models are seen in Fig. S16. Their confidence scores and PAE plots are seen together with those of the high-confidence models in Table S4 and Fig. S14, respectively.

The interactions between residues in the high-confidence models were visualized (Fig. S17-S21) to identify amino acids potentially involved in the binding between Lhca and FNR proteins. Sequence alignments of these proteins across the detected organisms were then conducted, highlighting conserved residues that are likely central to these interactions. Within the Lhca proteins, a small set of residues emerges as strongly conserved across all identified species. These residues cluster predominantly in the N-terminal region (Fig. 5C) and primarily form hydrophobic contacts with the FNR proteins, underscoring the central role of the N-terminal helix in the predicted interactions. Additional conserved N-terminal residues display lineage-specific patterns, mainly distinguishing *Chlamydomonas* species from *Monoraphidium* species together with *R. subcapitata*, reflecting their evolutionary relationships, as *Monoraphidium* and *Raphidocelis* belong to the Selenastraceae family^51^, while *Chlamydomonas* belongs to the Chlamydomonadaceae family^52^. Several predicted interaction sites are species-specific and are shown in the full Lhca sequence alignment (Fig. S22). FNR sequence alignments (Fig. S23) similarly reveal a group of widely conserved residues and lineage-specific divergence within the predicted binding interface. Notably, all Lhca and FNR residues that are preserved across the analyzed taxa interact with one another, suggesting they are the main mediators of the interaction between these proteins.

As previously noted, FNR binds to thylakoid membranes via Tic62 and TROL in higher plants, prompting the question of whether their FNR-binding regions resemble those identified in green microalgae. To address this, the FNR-binding motifs of Tic62 and TROL were aligned with the N-terminal helices of the algal Lhca proteins identified in this work (Fig. S24A). This comparison reveals limited conservation, including a hydrophobic residue at position 10 and a positively charged residue at position 14 in *Chlamydomonas* and *R. subcapitata*, corresponding to a Lys at this position in the Tic62/TROL motif. Alignment to the full length Lhca proteins was also conducted (Fig. S24B), showing two amino acids of *Monoraphidium* aligning with the Tic62/TROL motif. Despite these few similarities, the FNR-binding sites of algal Lhca proteins compared to those of Tic62 and TROL appear largely distinct.

## Discussion

The binding of FNR to thylakoids membranes is well characterized in higher plants and cyanobacteria, where it occurs via specialized proteins and structural adaptations. Much less is known about this process in green microalgae and other organisms, especially under ambient conditions. Prior work from our lab showed that FNR directly binds to PSI-LHCI via the LHCI complex. Here, we successfully identified and characterized the binding interface, pinpointing the N-terminal helix of Lhca4 as the region interacting with FNR by integrating Cryo EM data with AF modeling and ITC validation. Structural models indicated that FNR and Fd are unlikely to directly interact while both are bound to PSI-LHCI, although the bound FNR appears to be competent for Fd interaction. An evolutionary analysis suggests that the FNR-Lhca4 interaction is largely conserved in other green microalgae and may extend to other photosynthetic lineages, representing a distinct mechanism of FNR association with thylakoid membranes, differing from those observed in higher plants and cyanobacteria.

### Mechanistic insights into FNR binding at Lhca4

Earlier work^32^ demonstrated that FNR interacts with PSI in an LHCI dependent manner. Consistently, we identified the Lhca4 N-terminal helix as an FNR-binding domain. ITC measurements corroborated this interaction, showing an exothermic profile and thermodynamic parameters similar to those of FNR binding the intact PSI-LHCI complex (Table 1), with enthalpy contributions dominating over entropy, indicating that both interactions are primarily driven by contacts formation, as those depicted in (Fig. 2B-C), rather than by entropy gains from de-solvation or conformational disorder. Notably, the K_D_ of the interaction with the Lhca4-derived peptide was about one sixth of the K_D_ obtained for the full PSI-LHCI, perhaps due to greater accessibility of the free peptide. The absence of additional binding events detected by ITC, which is sensitive to affinities up to 10^-2^ M^55^, suggests that any other contacts are extremely weak and are unlikely to play a major role in anchoring FNR to PSI-LHCI. In addition, the predicted number of displaced water molecules is comparable between the peptide only and the full complex measurements, with discrepancies likely reflecting the flexibility of the FNR and PSI-LHCI, while the AF model represents a single, static state, yielding a smaller interface. Altogether, these findings suggest that the interaction between FNR and the N-terminal helix of Lhca4 constitutes the principal binding event mediating FNR association with PSI-LHCI.

**Table 1.** Comparison of FNR binding to Lhca4-derived peptide and PSI-LHCI. Summary of the thermodynamic parameters measured by ITC for FNR interaction with a peptide corresponding to the N-terminus of Lhca4 (this study) and with PSI-LHCI^32^. Shown are the binding stoichiometry (N), dissociation constant (K_D_), enthalpy change (ΔH), Gibbs free energy (ΔG), and entropy contribution (-TΔS); with standard deviations (n=4).

|  | FNR and Lhca4-derived peptide (Current Work) | FNR and PSI-LHCI (Marco et al. 2019) |
| --- | --- | --- |
| N | $1.07 \pm 0.03$ | $0.94 \pm 0.02$ |
| $K_D$ [nM] | $125.75 \pm 58.17$ | $780 \pm 110$ |
| $\Delta H$ [kcal/mol] | $-23.23 \pm 0.97$ | $-20.7 \pm 0.8$ |
| $\Delta G$ [kcal/mol] | $-9.4 \pm 0.11$ | $-8.3$ |
| $-T\Delta S$ [kcal/mol] | $13.83 \pm 0.46$ | $12.4$ |

Integration of the Cryo EM data with AF modeling suggests that FNR and Fd bind PSI-LHCI at distinct sites. The spatial separation between the two proteins is too great to support direct efficient electron transfer (Fig. 4). Nonetheless, the Lhca4 N-terminal helix binds FNR opposite to its FAD cofactor, allowing reduced Fd released from PSI to potentially engage PSI-bound FNR. Moreover, we cannot exclude the possibility that transient rearrangements of FNR might occasionally bring the two proteins into a catalytically competent orientation. Overall, electron transfer between FNR and Fd appears to occur sequentially rather than through simultaneous binding.

### Evolutionary conservation of the FNR binding helix

A bioinformatic search across cyanobacteria and all eukaryotic lineages showed that the FNR-Lhca4 interaction observed in *C. reinhardtii* is conserved in several other green microalgae, as indicated by high-confidence predictions of their FNR-Lhca complexes (Fig. 5). Considering models that satisfied all criteria, except for a sufficient ipTM score, enabled the detection of similar interactions across additional algal taxa. These results highlight the conservation of FNR-Lhca4 interactions in green microalgae, without precluding other potential binding mechanisms in certain species. Extending our analysis to low-confidence models also revealed that *Emiliania huxleyi*, a coccolithophore, may possess an FNR-interacting Lhca N-terminal helix. This observation is consistent with the emergence of analogous FNR membrane-association mechanisms across different phyla. For example, an additional membrane tethering mechanism may operate in maize and wheat FNR isoforms which were shown to bind thylakoid membranes by their N-terminus^56,57^, reminiscent of the prevalent cyanobacterial mechanism. Consistent with our demonstration of direct FNR binding to PSI, similar associations have been reported in barley^58^ and *Nannochloropsis oceanica*^59^. However, in these species, FNR interacts with PSI core subunits rather than with light harvesting proteins. Therefore, the observed FNR-Lhca interactions are likely not restricted to green microalgae and could represent a broader strategy conserved across other photosynthetic lineages.

In this work, we uncover a previously unrecognized mode of FNR association with thylakoid membranes in *C. reinhardtii*, likely conserved across other green microalgae and possibly additional organisms, mediated by the N-terminal helix of Lhca4. This mechanism is distinct from FNR recruitment in higher plants, which relies on Tic62 and TROL, whose FNR-binding motifs differ substantially from those described here (Fig. S24), and from cyanobacteria, where FNR binds PBSs via an intrinsic extension. We propose that direct interaction of FNR with PSI-LHCI via Lhca4 is responsible for the PSI-associated FNR observed in the absence of PGR5 and/or PGRL1^29^. In the suggested model (Fig. 6), the N-terminal helix of Lhca4 functions as a transient docking site, positioned at a distance that avoids steric interference with Fd. Under anaerobic conditions, stabilization of FNR at PSI by PGR5 and PGRL1 accompanied by dissociation of Lhca2 and Lhca9^31^ further promotes CEF at the expanse of LEF, consistent with enhanced CEF observed when FNR is fused to PSI^62^. Altogether, these findings reveal a novel mechanism for FNR association with thylakoid membranes in green microalgae and phylogenetically distant species such as *Emiliania huxleyi*, advancing our understanding of photosynthetic electron flow regulation.

**Fig. 6.**
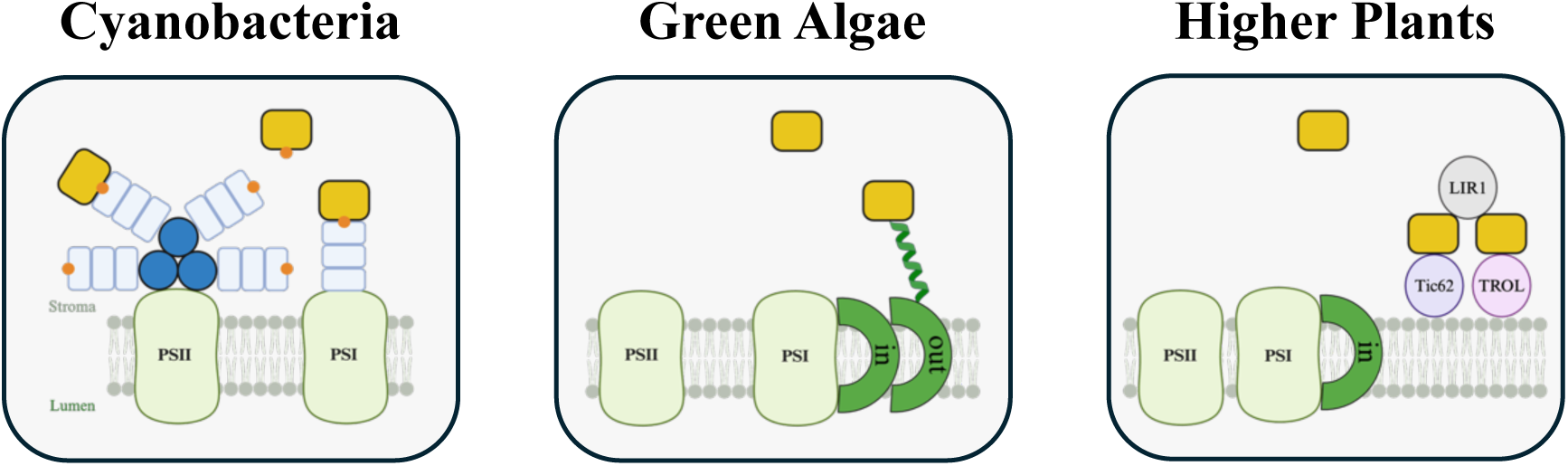
FNR binding to thylakoid membranes across photosynthetic organisms. Schematic comparison of the mechanisms by which FNR (yellow squares) associates with thylakoid membranes. In cyanobacteria, FNR binds to PBSs found both on PSII and PSI through its CpcD-like domain (orange dots)^60^. In higher plants, FNR is recruited by Tic62 and TROL, with LIR1 stabilizing these interactions^61^. Our data suggest that in green algae, FNR binds the N-terminal helix of Lhca4, which is located in the outer LHCI belt (“out”), present only in green algae, whereas “in” refers to the inner LHCI belt, conserved both in green algae and higher plants. Figure created with BioRender.

## Methods

### Proteins Purification

FNR, Fd and SOD of *C. reinhardtii* were heterologously expressed in *E. coli* and purified as previously described^63,64^. PSI-LHCI was isolated from the JVD1B *C. reinhardtii* strain according to^64^.

### Cryo EM

PSI-LHCI complexes were incubated (I) with FNR and Fd at a 1:10:10 molar ratio at pH 7.5 and (II) pH 6.5, (III) with SOD or (IV) with FNR alone at a 1:10 molar ratio (Fig. S1-S4, Table S1). Samples were applied to glow-discharge holey carbon grids and vitrified using Leica EM GP, with blotting performed for 2.5 sec at 4°C and 100% humidity. Cryo EM data were acquired on a 300 kV Titan Krios G3 electron microscope (Thermo Fisher Scientific) equipped with a Gatan BioQuantum energy filler and a K3 Summit direct electron detector (Ametek), operated using EPU 1.9 software. Movies were recorded in counting mode at ×105,000 magnification, corresponding to a calibrated pixel size of 0.85 Å, with a total electron dose of 42 e^-^ Å^-2^ and a defocus range of 0.7-2.5 μm. A total of 8,838, 7,466, 10,284 and 10,564 micrographs were collected for datasets (I-IV), respectively. Image processing was carried out in cryoSPARC^65^. Movies were subjected to motion correction, followed by contrast transfer function (CTF) estimation. Particle picking, classification and refinement procedures are described in Fig. S1-5. From the four datasets (I-IV), 203,662, 130,754, 138,168 and 99,327 high-quality particles, respectively, were selected and refined to final resolution of 2.2, 2.29, 2.92 and 2.5 Å. Given the apparent flexibility of FNR, focused classification was performed using signal subtraction with a mask encompassing the FNR region. Classification into four, six or ten classes was tested (Fig S5).

### AlphaFold

All models were generated using AlphaFold 3 via the AlphaFold server^66^ under its default configuration. Unpaired multiple sequence alignments (MSAs) for individual chains were obtained from UniRef90, MGnify and Small BFD; while paired MSAs for predicting inter-chain interactions were sourced from UniProt. PDB templates were restricted to structures released before September 20, 2021. The number of recycles was automatically set to 10, and no restraints were applied. For each prediction, only the top-ranked model was analyzed.

### AF modeling of FNR and Lhca proteins of *C. reinhardtii*

The mature FNR sequence was obtained from UniProt (Q9S9E0). Lhca1, 4, 6 and 5 sequences were retrieved from Phytozome (*C. reinhardtii* CC-4532 v6.1 genome) and their mature form was determined based on a previous study that performed N-terminal sequencing of the Lhca proteins of *C. reinhardtii*^33^.

### ITC

#### Background

Isothermal titration calorimetry (ITC) is a well-established technique for quantifying biomolecular interactions through measurement of the heat released or absorbed upon binding^67,68^. The instrument contains a sample cell, loaded with the macromolecule of interest; and a reference cell, filled with the solvent of the sample to correct for baseline and instrumental thermal fluctuations^69^. During an experiment, successive amounts of titrant are injected into the sample cell via a syringe, and the heat released or absorbed by the binding events leads to a deviation of the cell temperature. In response, power is applied to restore the set temperature, and is recorded over the course of the experiment, resulting in peaks from which thermodynamic parameters are derived. As binding progresses, the heat peaks gradually decrease, reflecting saturation of the molecules in the cell^68^. Downward peaks indicate an exothermic interaction, where heat is released and the system responds by cooling. Conversely, upward peaks correspond to an endothermic interaction, in which heat is absorbed, and the system compensates by warming. For each titrant addition, the area of the resulting peak is computed to determine the heat change^69^. The binding stoichiometry (n), dissociation constant (K_D_; reciprocal to the association constant K_a_) and enthalpy change (ΔH) are obtained by employing nonlinear regression to fit the data to a binding model of the interaction, such as one that assumes that a single ligand binds to a single site on the macromolecule^70,71^. The free energy (ΔG) is derived from the dissociation constant^71^:

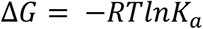

The entropy change is then calculated using the relation:

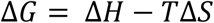

#### Experimental setup

FNR and the synthesized peptides were diluted to a final concentration of 10 μM and 100 μM, respectively, in 50 mM Tris-HCl pH 8. ITC experiments were performed using MicroCal PEAQ-ITC instrument (Malvern) at 25°C. The reference power was set to 5 μcal/second, the feedback to “high”, and the stirring speed to 750 RPM. Thirteen injections were performed, beginning with a 0.4 μL injection (0.6 seconds duration, 60 seconds spacing) to equilibrate the system and establish a stable baseline; this injection was excluded from analysis. Twelve subsequent 3 μL injections (6 seconds duration, 180 seconds spacing) were used for binding measurements. The resulting data were analyzed in the MicroCal PEAQ-ITC analysis software (Malvern), where they were fitted using a one set of sites model to obtain thermodynamic parameters.

### Peptides Synthesis and Preparation

The synthetic peptides utilized in the ITC experiments were synthesized by PEPperPRINT (Heidelberg, Germany) with ≥ 95% purity. The sequence of the native peptide is SAVPENVKEAREWIDAWKSKSGG. The sequence of the mutated peptide is SAVPRNVKHARHWIKAWKSKSGG.

### Circular dichroism (CD) spectroscopy

Peptides were initially dissolved in ultra-pure water to create a 10 mM stock solution, which was then diluted with 5 mM Tris-HCl buffer at pH 8 to achieve a final concertation of 1 mg/mL. CD measurements were performed using a Chirascan spectrometer (Applied Photophysics, Leatherhead, UK) equipped with a Peltier temperature control unit set to 25°C. Spectra were acquired in 0.1 mm path length quartz cuvettes (Hellma Analytics) across 190-260 nm with a step size of 1 nm, a spectral bandwidth of 1 nm, and an integration time of 1 sec per point. The averaged spectrum of three measurements of each peptide is presented. A buffer-only blank was recorded at the same conditions and subtracted from the peptide measurements. The resulting data were processed using Pro-Data Viewer software (Applied Photophysics).

### Bioinformatic search for orthologs

To evaluate whether the interaction between FNR and Lhca4 observed in *C. reinhardtii* is conserved, we performed BLASTP searches^72^ using Lhca4 amino acid sequence devoid of its signal peptide across the following taxonomic groups: *Cyanobacteria*, SAR (*Stramenopiles, Alveolata*, and *Rhizaria*), *Hacrobia, Unikonts, Excavata, Chlorophyta, Glaucophyta, Rhodophyta*, *Embryophyta*, and *Charophyta*. Taxonomic identifiers for each group are provided in Table S2. Where no single taxon ID encompassed an entire group, relevant subgroups were included. These groups and subgroups were selected based on established phylogenetic trees, which depict the relationships among all eukaryotes^53^, and more specifically, among major lineages of the plant kingdom^54^. Although the subgroups of *Unikonts* (*e.g, Opisthokonta*, which comprises animals and fungi) are, to the best of current knowledge, non-photosynthetic or predominantly heterotrophic^73^, we nevertheless included them in our BLASTP search to exclude any unexpected homologs.

Sequences were selected if they aligned with the first 21 amino acids of *C. reinhardtii* Lhca4. Their signal peptides were predicted using TargetP (organism group: plant)^74^. If the prediction was unsuccessful, SignalP (organism group: eukarya)^75^ was employed. The TargetP and SignalP prediction are shown in Fig. S6. Sequences for which the signal peptide could not be confidently determined were kept in their entirety, under the assumption that they are already processed, whereas the predicted signal peptides were omitted from sequences prior to submission to AF. Only proteins with a predicted global fold confidence (pTM score) greater than 0.5 and an N-terminal region predicted to form one α-helix facing towards FNR, as observed in *C. reinhardtii*, were retained. In addition, only Lhca proteins that match the canonical structure of Lhca proteins (as described in the Results section) were kept.

Next, we sought to identify FNR homologs in each organism by performing BLASTP search using the *C. reinhardtii* FNR sequence (UniProt Q9S9E0) against each species (their taxonomic identifiers are provided in Table S3). Signal peptides were determined as described above, predictions are shown in (Fig. S7). The resulting Lhca and FNR proteins were then modeled together by AF to predict their interaction.

### Visualization and interaction analysis of structural models

Generated AF models and experimentally resolved Cryo EM structures were visualized and inspected using ChimeraX^76–78^. For the FNR-Lhca4 AF model, interacting residues were identified based on spatial proximity and chemical properties. Hydrophobic contacts were defined as interactions between hydrophobic residues (Ala, Val, Leu, Ile, Met, Phe, Trp and Pro) with carbon-carbon distances of less than 5 Å^79^. Hydrogen bonds and salt bridges were detected using ChimeraX *h-bonds* tool, with hydrogen bonds identified under default settings and salt bridges identified using the “salt bridges only” option. A distance cutoff of 3.5 Å for hydrogen bonds^80^ and 4 Å for salt bridges^81^ with a 20° angle tolerance were applied. The interaction interface was calculated using the “measure buriedarea” command.

### Sequence alignments

MAFFT (version 7) was used to perform alignment of sequences using its default settings, except the output order was set to “same as input”^82^. The alignments were visualized using Jalview (version 2.11.5.0)^83^.

## Acknowledgments

We gratefully acknowledge Dr. Andreas Naschberger and Prof. Alexey Amunts (SciLifeLab, Stockholm, Sweden) for their exceptional support and expertise in Cryo-EM data acquisition. Their generous allocation of microscope time and assistance in collecting high-quality Cryo-EM datasets were essential to the success of this study. This work was supported by the Israel Science Foundation (Grant No. 941/22). Y.M is supported by the U.S. National Science Foundation under Grant No. 2145562.

## Author Contributions

S.A. designed and carried out the experiments, performed protein purification and ITC measurements, analyzed the data, and wrote the manuscript. P.M. purified PSI, Fd and FNR. T.E. purified PSI. O.B.Z. purified SOD. Y.M. co-designed the study, led data analysis, and co-wrote the manuscript. I.Y. conceived and supervised the study, secured funding, and co-wrote the manuscript.

