## Supplemental figures for "A Conserved Mechanism for Positioning Ferredoxin–NADP⁺ Reductase at Photosystem I in Green Algae"

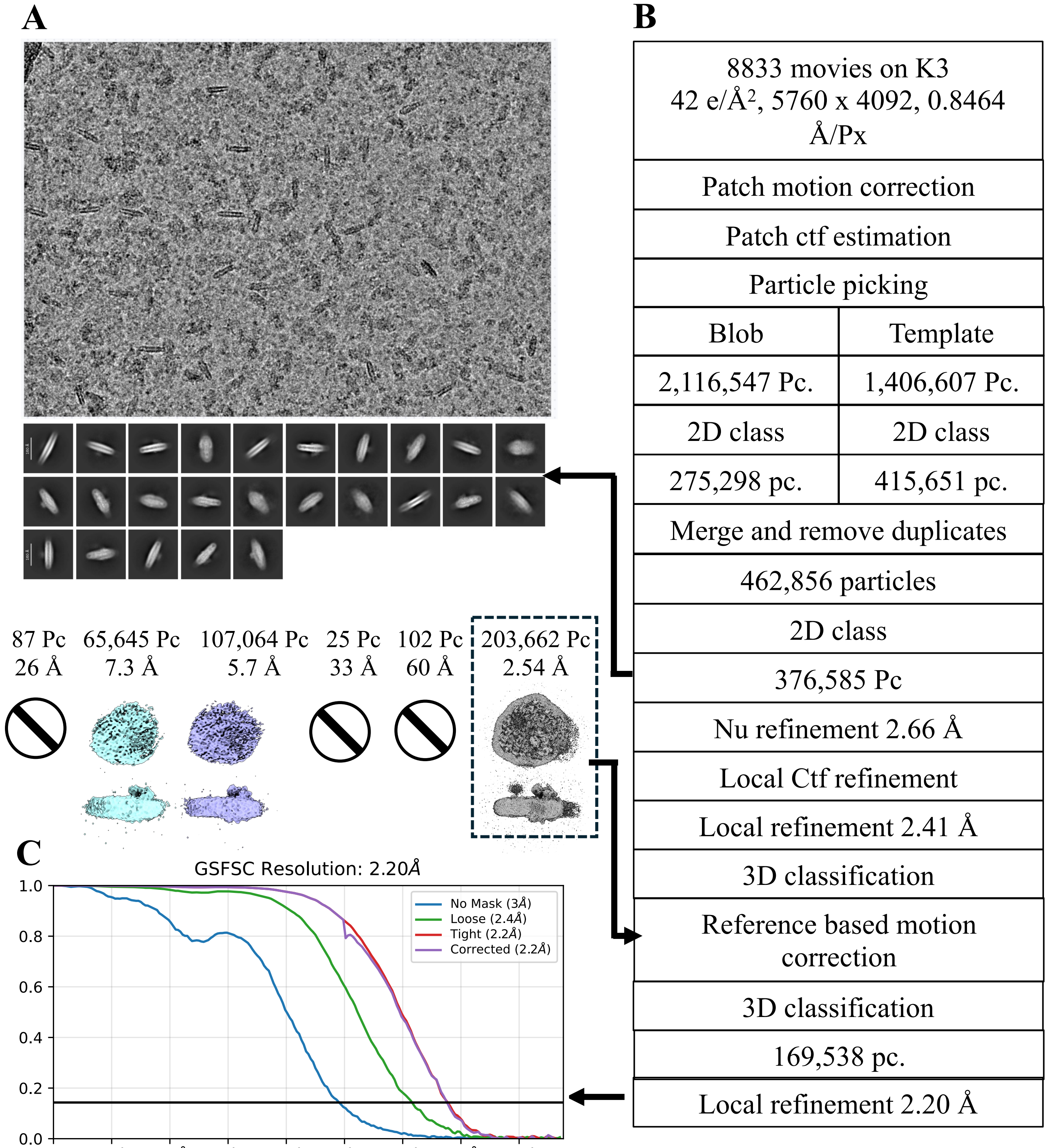


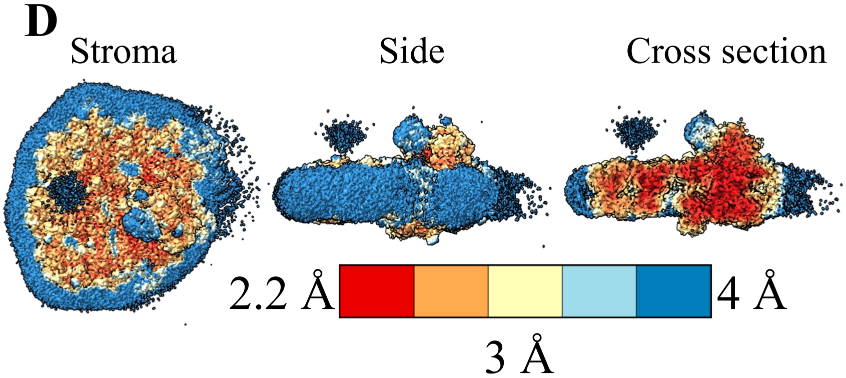


EMD-75304, PDB ID 10NJ

**Fig. S1. Cryo EM data processing of PSI-LHCI in complex with FNR and Fd at pH 7.5. (A)** Sample micrograph after motion correction. **(B)** Different stages of data process, all carried out using CryoSPARC. **(C)** FSC curves after referenced-based motion correction. **(D)** Local resolution after reference-based motion correction. (assigning peripheral regions to high resolution is known artefact and does not represent high resolution data).

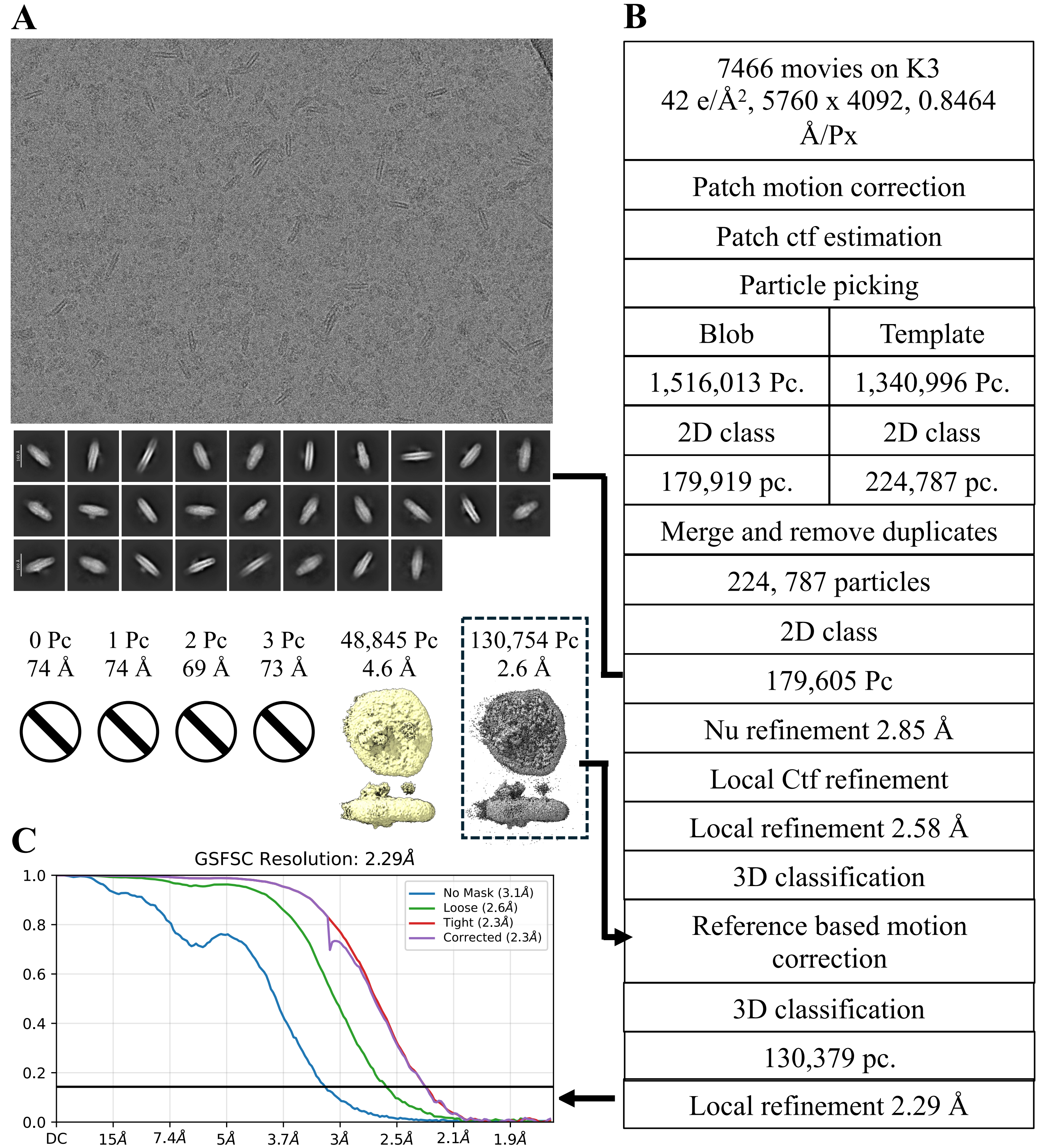

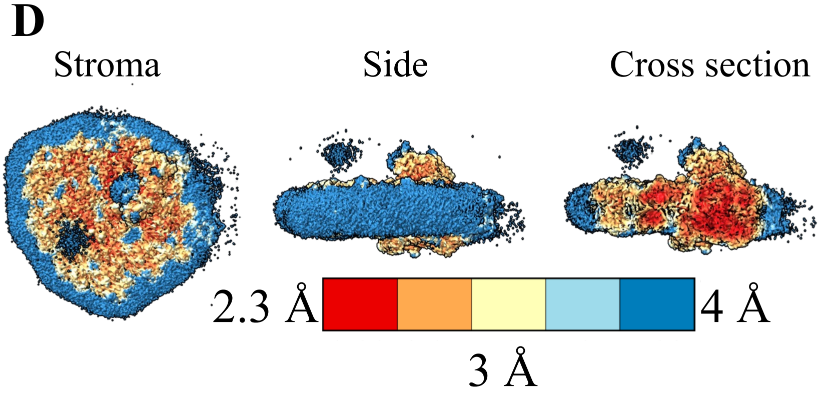


**Fig. S2. Cryo EM data processing of PSI-LHCI in complex with FNR and Fd at pH 6.5. (A)** Sample micrograph after motion correction. **(B)** Different stages of data process, all carried out using CryoSPARC. **(C)** FSC curves after referenced-based motion correction. **(D)** Local resolution after reference-based motion correction.

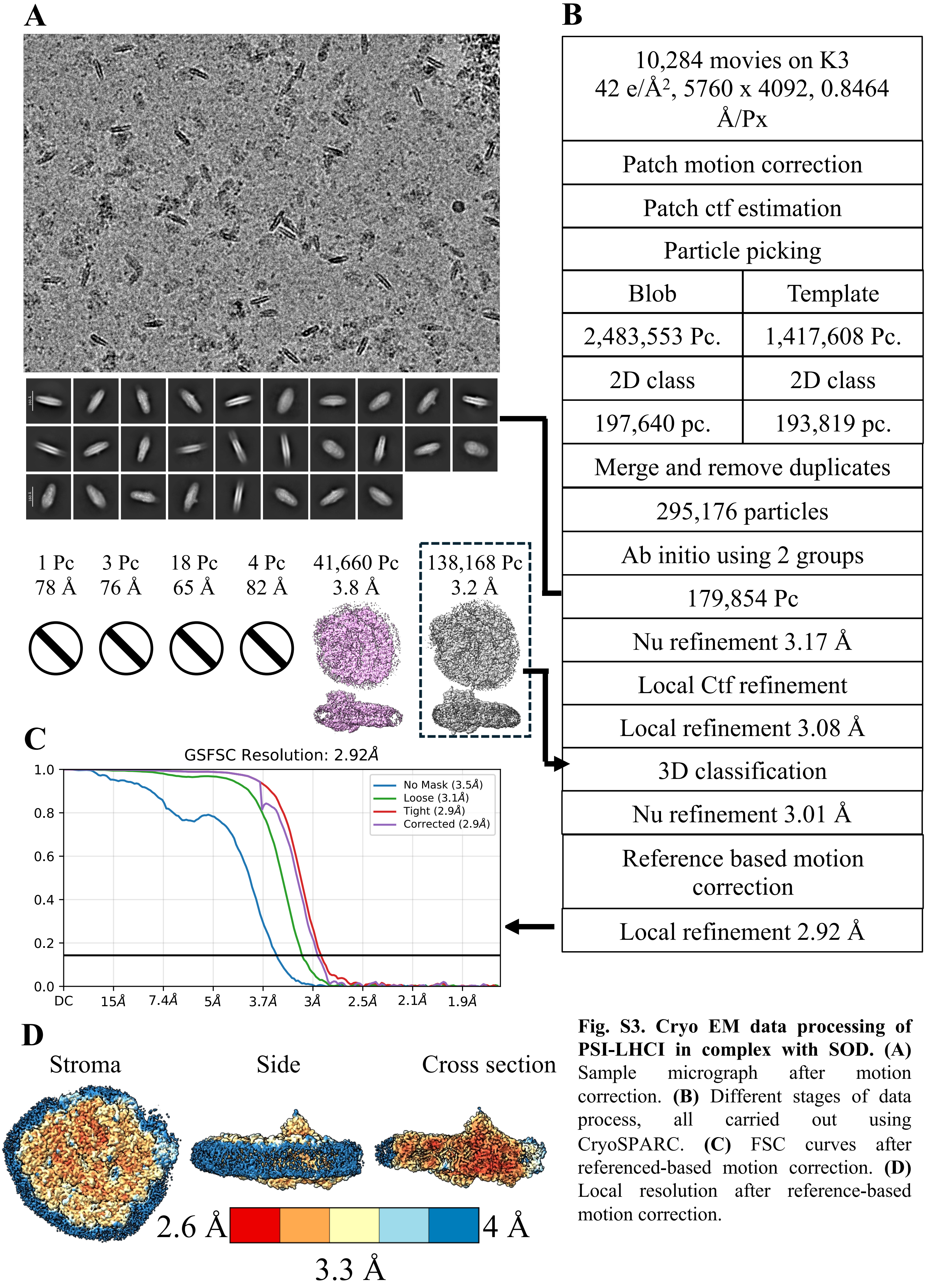


**Fig. S3. Cryo EM data processing of PSI-LHCI in complex with SOD. (A)** Sample micrograph after motion correction. **(B)** Different stages of data process, all carried out using CryoSPARC. **(C)** FSC curves after referenced-based motion correction. **(D)** Local resolution after reference-based motion correction.

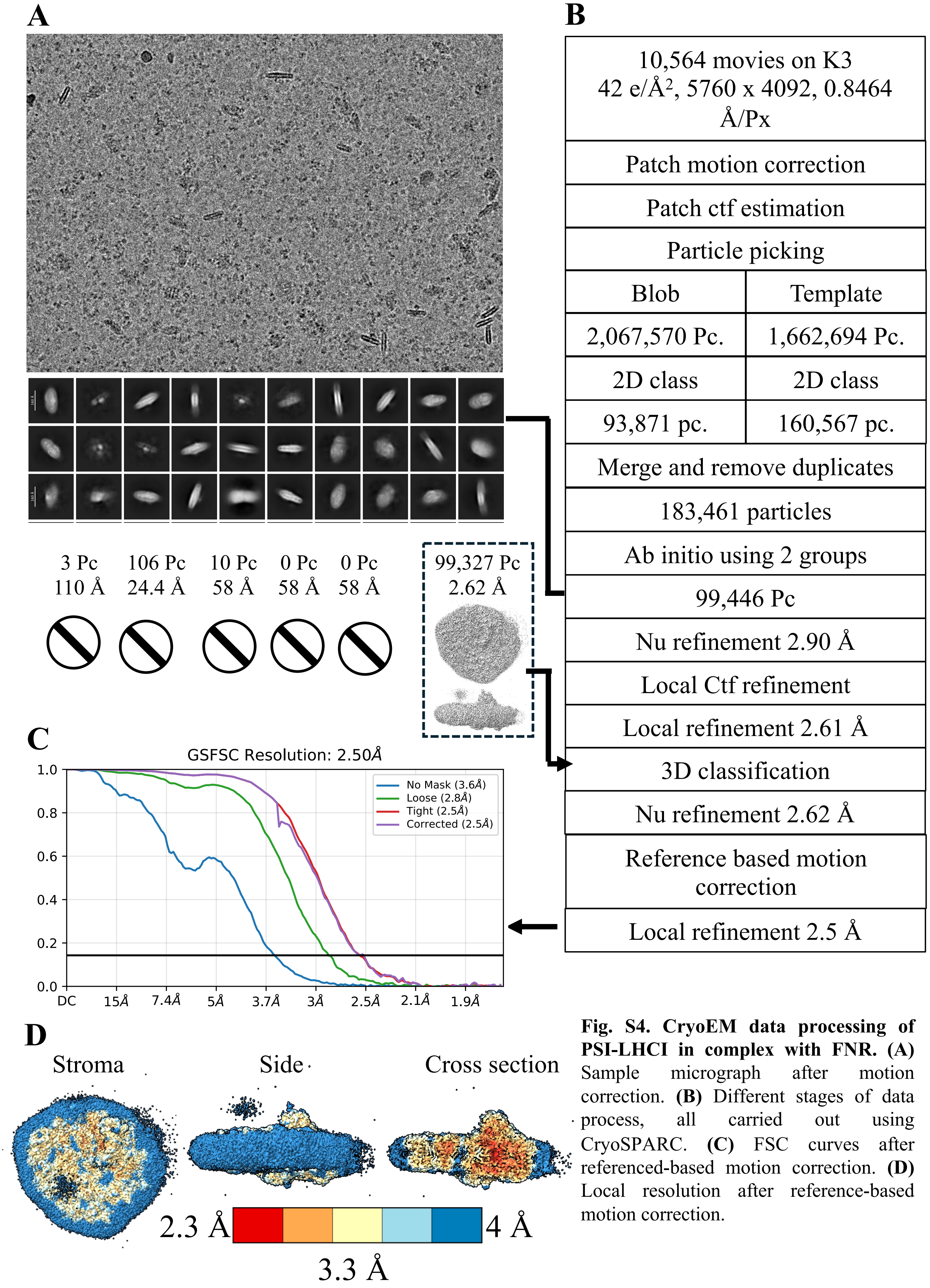


**Fig. S4. CryoEM data processing of PSI-LHCI in complex with FNR. (A)** Sample micrograph after motion correction. **(B)** Different stages of data process, all carried out using CryoSPARC. **(C)** FSC curves after referenced-based motion correction. **(D)** Local resolution after reference-based motion correction.

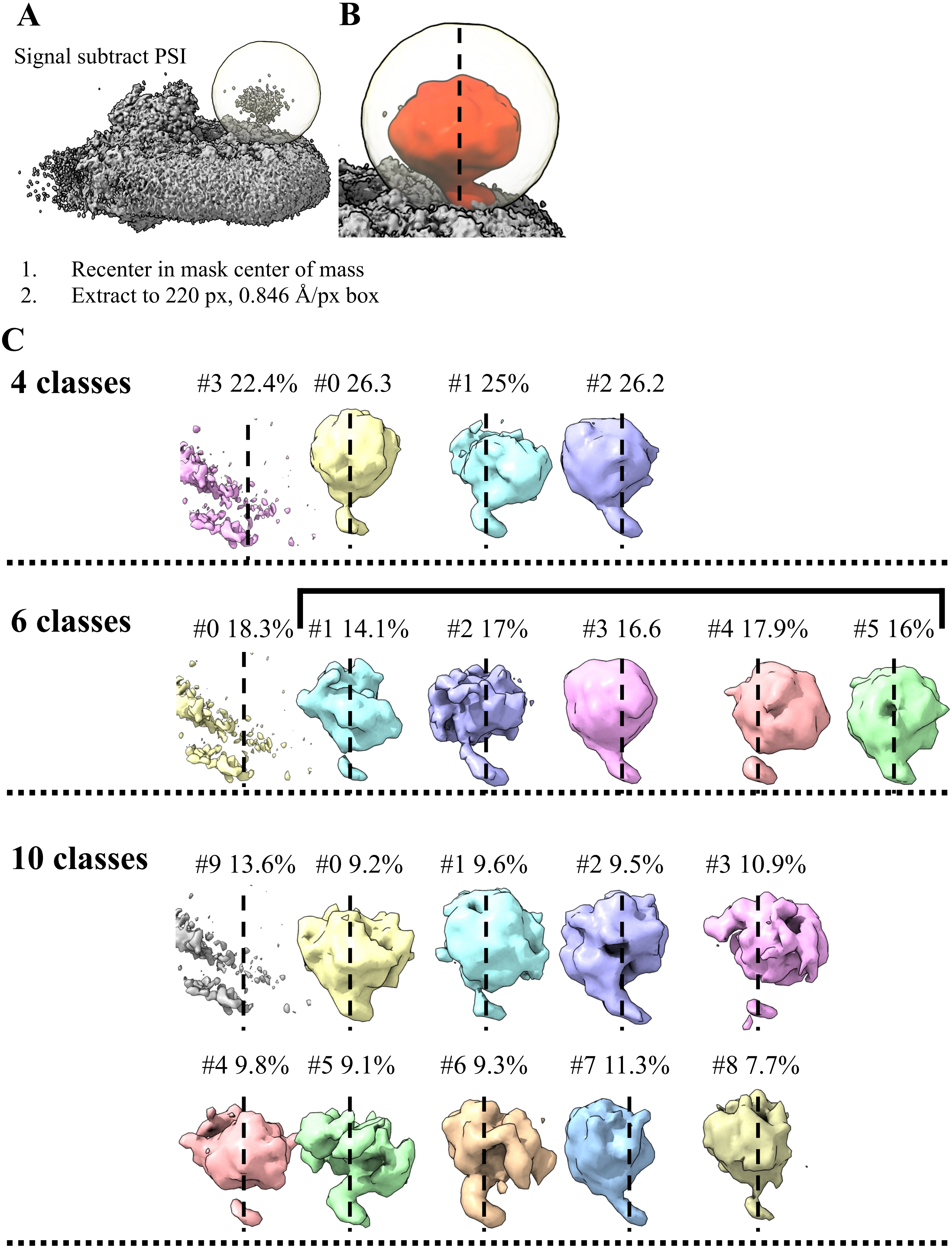


**Fig. S5. Cryo EM data processing – FNR classification and signal subtraction**. **(A)** The diffuse signal associated with FNR presence surrounded by the mask used for signal subtraction. **(B)** The same volume after subtracting the PSI signal. The original PSI-LHCI map is also shown in grey. **(C)** Three 3D classification jobs using different numbers of classes, showing the presence of one "noise" and additional diffuse classes at various orientations around the center of the volume (marked with a dashed line).

### *C. incerta*

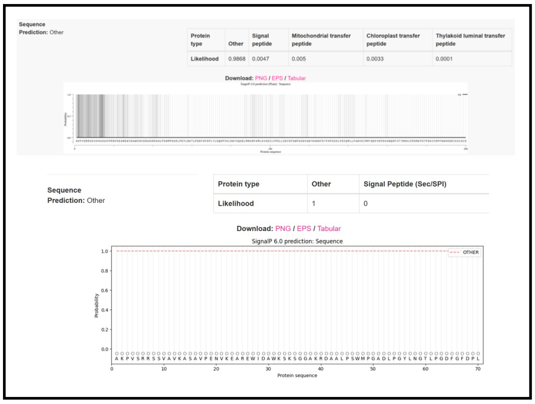


### *G. pectorale*

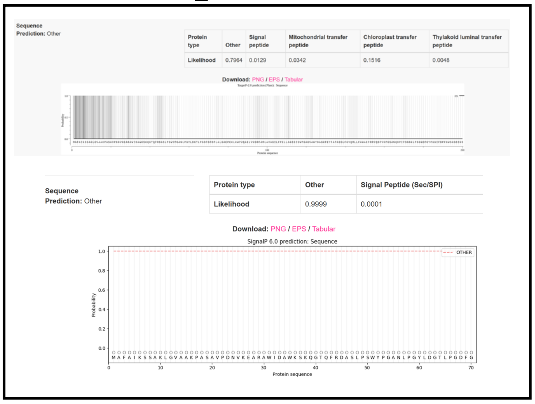


### *M. neglectum*

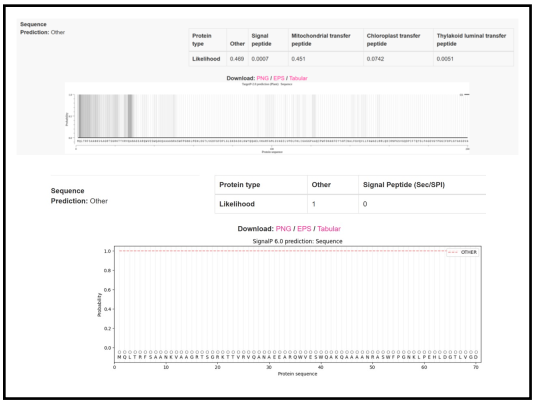


### *S. sp.* PABB004

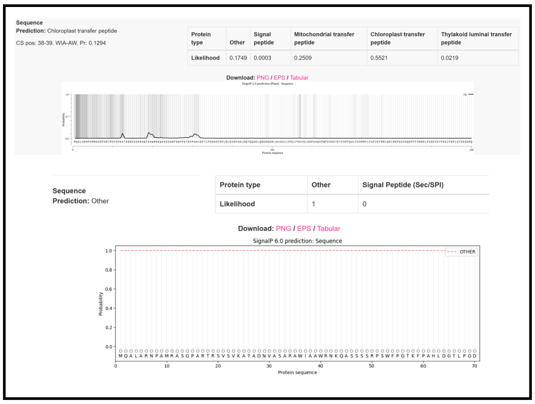


### *C. sp.* UWO 241

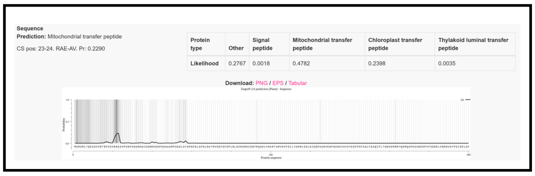


### *V. c. f. nagariensis*

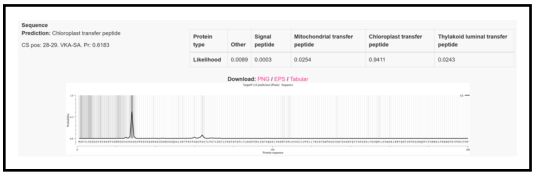


### *M. minutum*

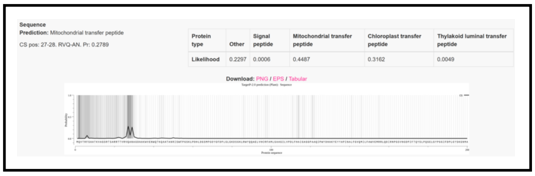


### *E. huxleyi* CCMP 1516

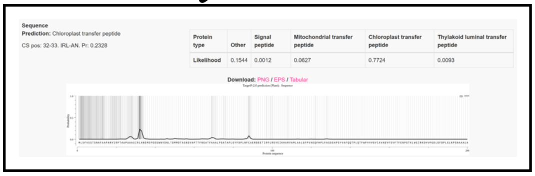


### *R. subcapitata*

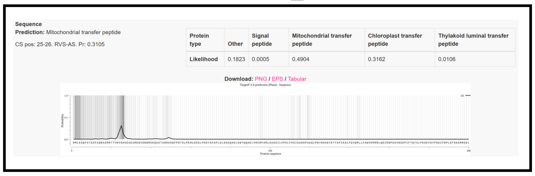


**Figure S6. Signal peptide predictions for Lhca4-homologs.**
Lhca proteins with an N-terminal region homologous to *C. reinhardtii* Lhca4 were analyzed using TargetP and/or SignalP to identify their transit peptides.

### *C. incerta*

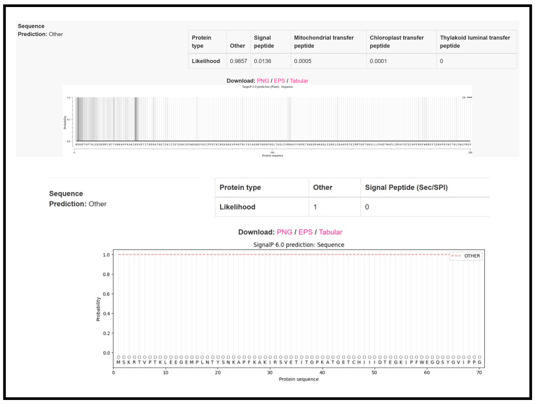


### *V. c. f. nagariensis*

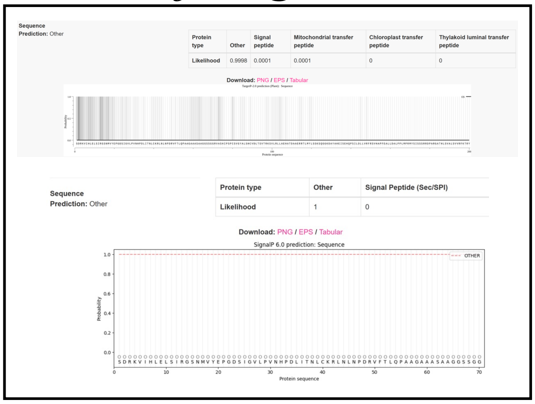


### *M. neglectum*

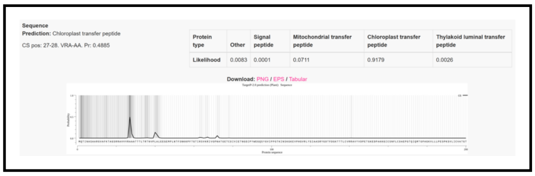


### *G. pectorale*

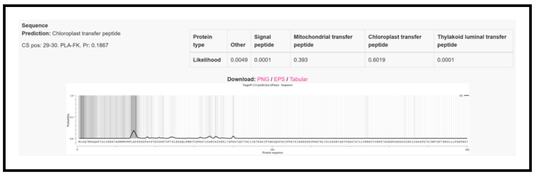


### *C. sp.* UWO 241

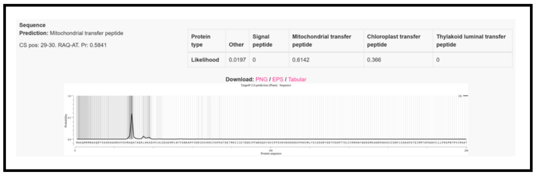


### *S. sp.* PABB004

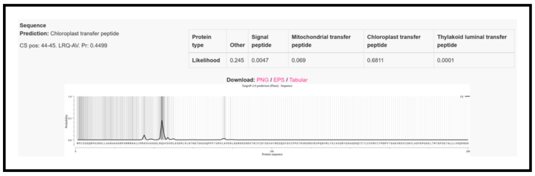


### *M. minutum*

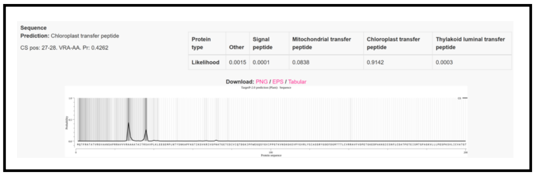


### *E. huxleyi* CCMP 1516

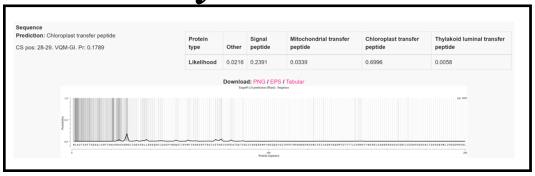


### *R. subcapitata*

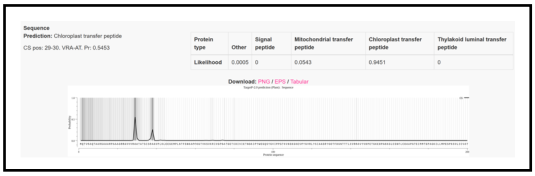


**Fig. S7. Signal peptide predictions of identified FNR proteins.**
FNR proteins from organisms possessing a Lhca protein with an N-terminal region homologous to *C. reinhardtii* Lhca4 were analyzed using TargetP and/or SignalP to identify their transit peptides.

A B

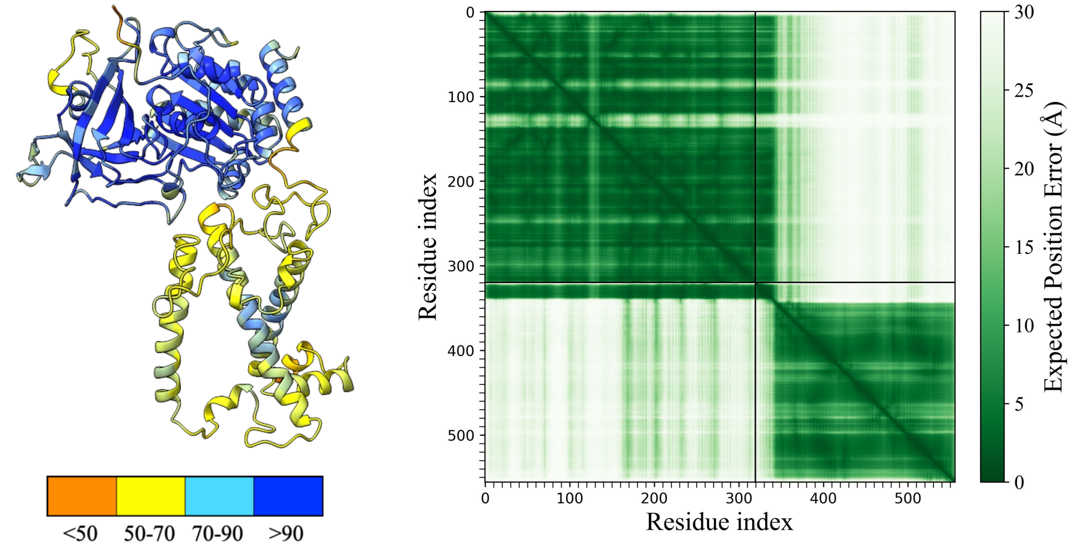


C

| Mean pLDDT | Helix Mean pLDDT | ipTM | pTM |
|---|---|---|---|
| 86.85 | 92.23 | 0.87 | 0.63 |

**Figure S8. Confidence parameters of *C. reinhardtii* FNR and Lhca4 AlphaFold model.** **(A)** The predicted structure colored by per-residue pLDDT scores: > 90 (blue), 70-90 (cyan), 50-70 (yellow) and <50 (orange). Higher scores indicate greater confidence in the local geometry of each residue **(B)** PAE plot of this model. As indicated by the bar, darker green corresponds to higher certainty in the relative positioning of the residues. The two protein chains (FNR and Lhca4, respectively) are concatenated and shown together. **(C)** mean pLDDT scores of the full model and of the Lhca4 N-terminal helix, together with the inter-chain TM (ipTM) and overall pTM scores.

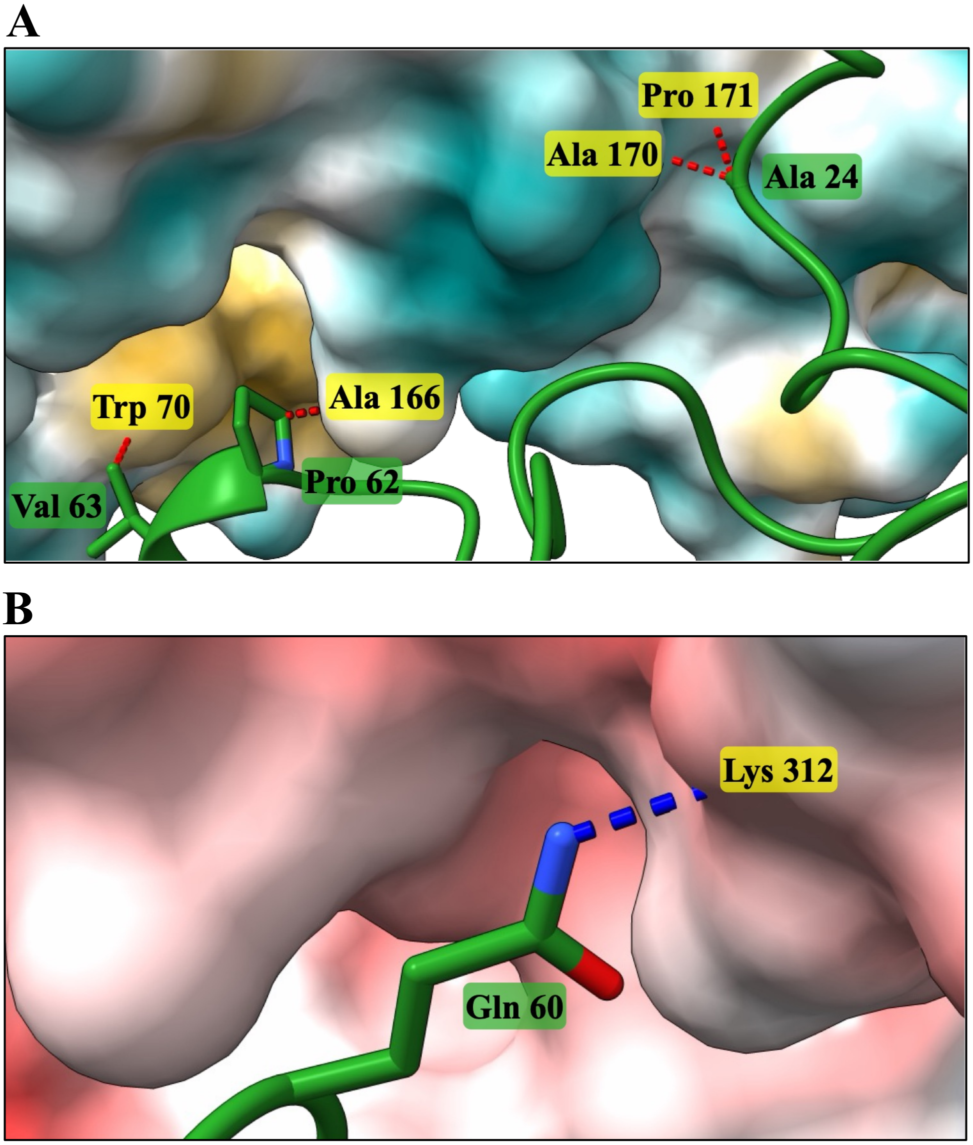


**Figure S9. Predicted FNR-Lhca4 interactions outside the N-terminal helix in *C. reinhardtii.***

Only the shortest distances between residue pairs are displayed, with FNR residues in yellow and Lhca residues in green. **(A)** Hydrophobic contacts outside the N-terminal helix, all distances <4.5 Å. The FNR surface is colored by hydrophobicity, with cyan indicating hydrophilic (polar) regions and golden-brown indicating hydrophobic (apolar) regions. **(B)** A hydrogen bond is observed at 2.96 Å. FNR surface is colored by electrostatic potential, ranging from red (negative) to blue (positive). No salt bridges contributing to the FNR-Lhca4 interface are detected outside the N-terminal helix.

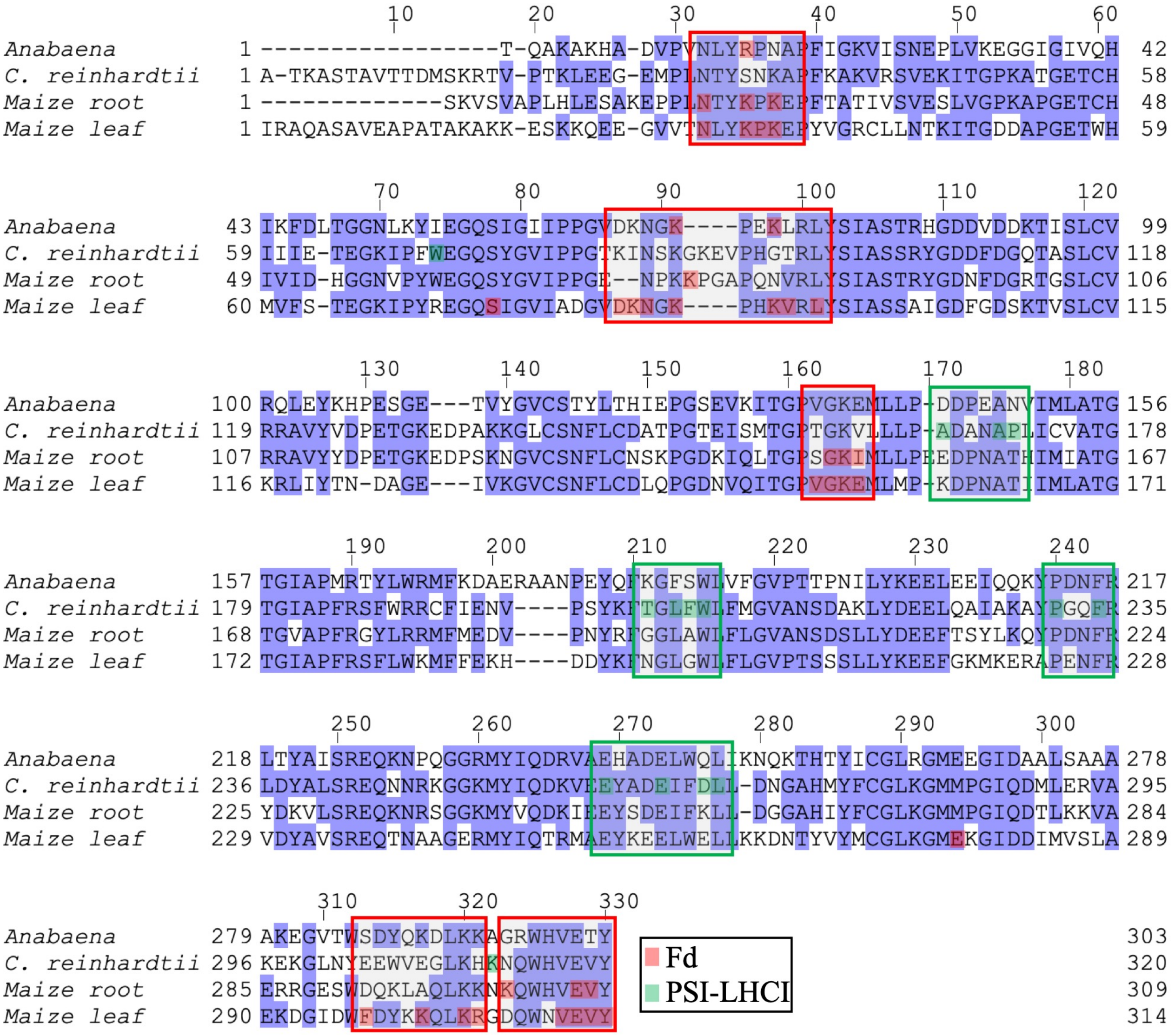


**Fig. S10. Residues involved in FNR interactions with Fd and PSI-LHCI.**
Sequence alignment of FNR from *Anabaena*, *C. reinhardtii*, and maize (leaf and root isoforms). Positions marked in red mediate FNR-Fd interactions in *Anabaena* or maize[37,39,44,45]. Positions marked in green contact Lhca4 in *C. reinhardtii*. Red or green frames highlight regions with predominantly FNR-Fd interacting residues or residues contacting Lhca4, respectively. Amino acids with $\geq$40% identity, defined as the fractions of identical residues among all non-gap positions, are colored in purple.

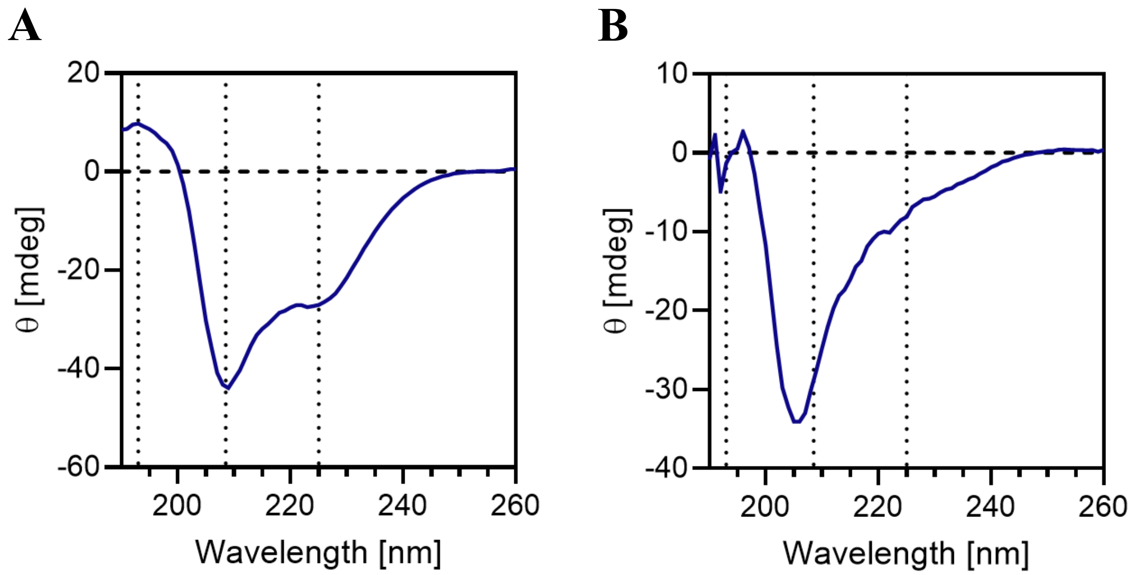


**Figure S11. Circular dichroism (CD) profile of the Lhca4-derived peptides.**
CD spectra of 1 mg of the native **(A)** or mutated **(B)** Lhca4-derived peptides utilized for ITC measurements in 5 mM Tris buffer pH 8.

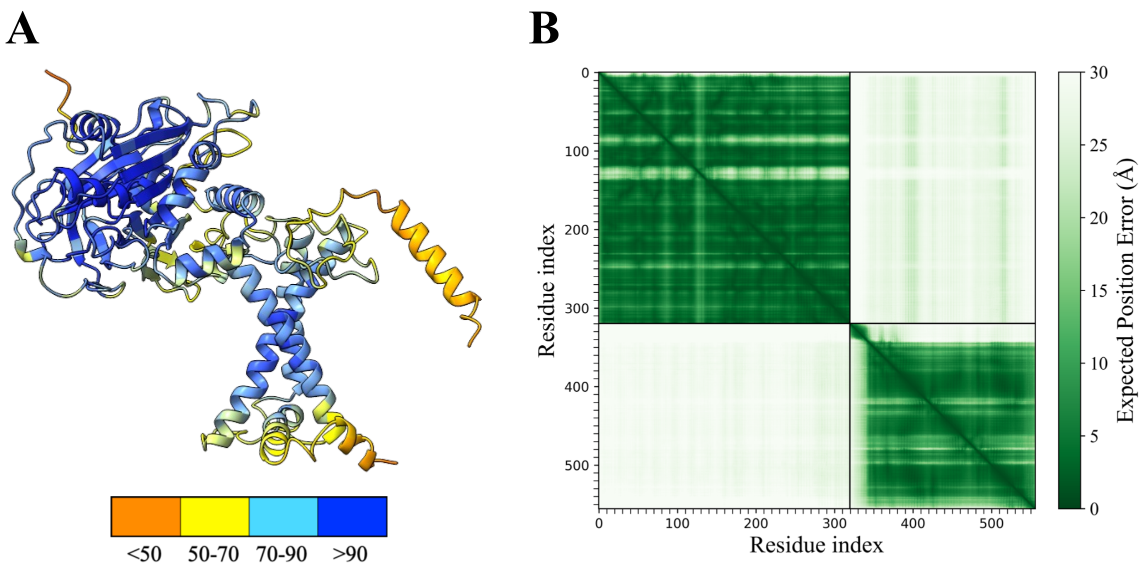


C

| Mean pLDDT | Helix Mean pLDDT | ipTM | pTM |
|---|---|---|---|
| 86.96 | 53.26 | 0.13 | 0.59 |

**Figure S12. Confidence parameters of *C. reinhardtii* FNR-Lhca4 AlphaFold model with a mutated Lhca4 helix.**

The Lhca4 N-terminal helix was modified by replacing all negatively charged residues with positive ones, causing it to point away from FNR. This helix was synthesized as a peptide and utilized in the ITC assays shown in Fig. 3. **(A)** The model colored by per-residue pLDDT scores, where higher scores indicate greater confidence in the local geometry. Residues with pLDDT > 90 are shown in blue, 70-90 in cyan, 50-70 in yellow and < 50 in orange. **(B)** PAE plot of this model, darker green indicating higher confidence in the relative positioning of the residues. FNR and Lhca4 are shown concatenated. **(C)** the mean pLDDT of the entire model, the mean pLDDT of the residues forming the mutated helix, as well as the inter-chain TM (ipTM) and overall pTM scores.

A

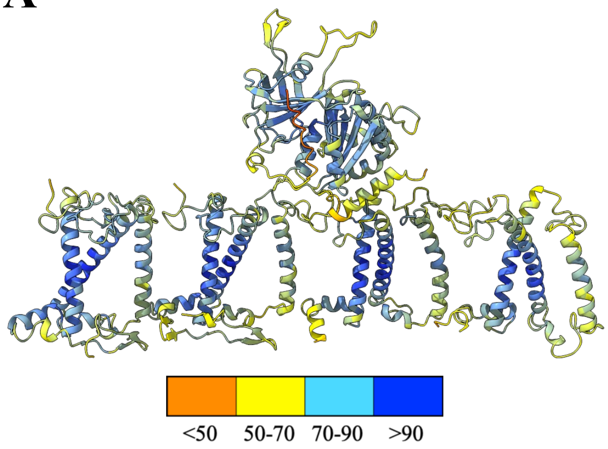


B

| | Mean pLDDT |
|---|---|
| All | 85.86 |
| FNR | 85.53 |
| Lhca4 | 84.75 |
| Lhca1 | 83.97 |
| Lhca6 | 86.71 |
| Lhca5 | 88.23 |

C

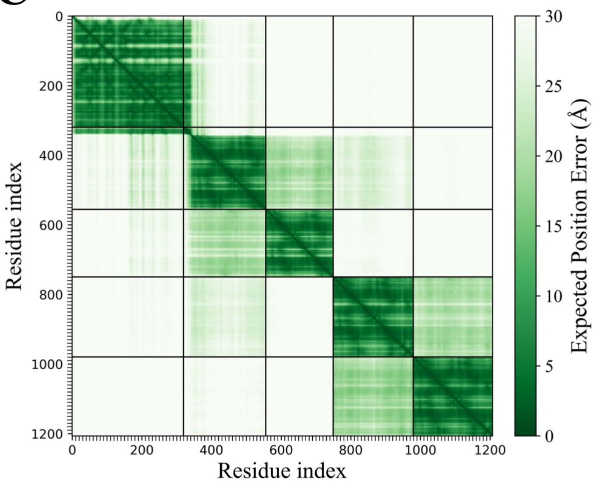


**Fig. S13. Confidence parameters of *C. reinhardtii* FNR and outer LHCI belt AlphaFold model.**
**(A)** The predicted structure colored by per-residue pLDDT scores: > 90 (blue), 70-90 (cyan), 50-70 (yellow) and <50 (orange). Left to right: Lhca5, Lhca6, Lhca4 and Lhca1. FNR is above Lhca4. Panel **(B)** lists the mean pLDDT scores of the model and of each chain. **(C)** PAE plot of the model. The inter-chain TM (ipTM) is 0.34 and the overall pTM score is 0.37.

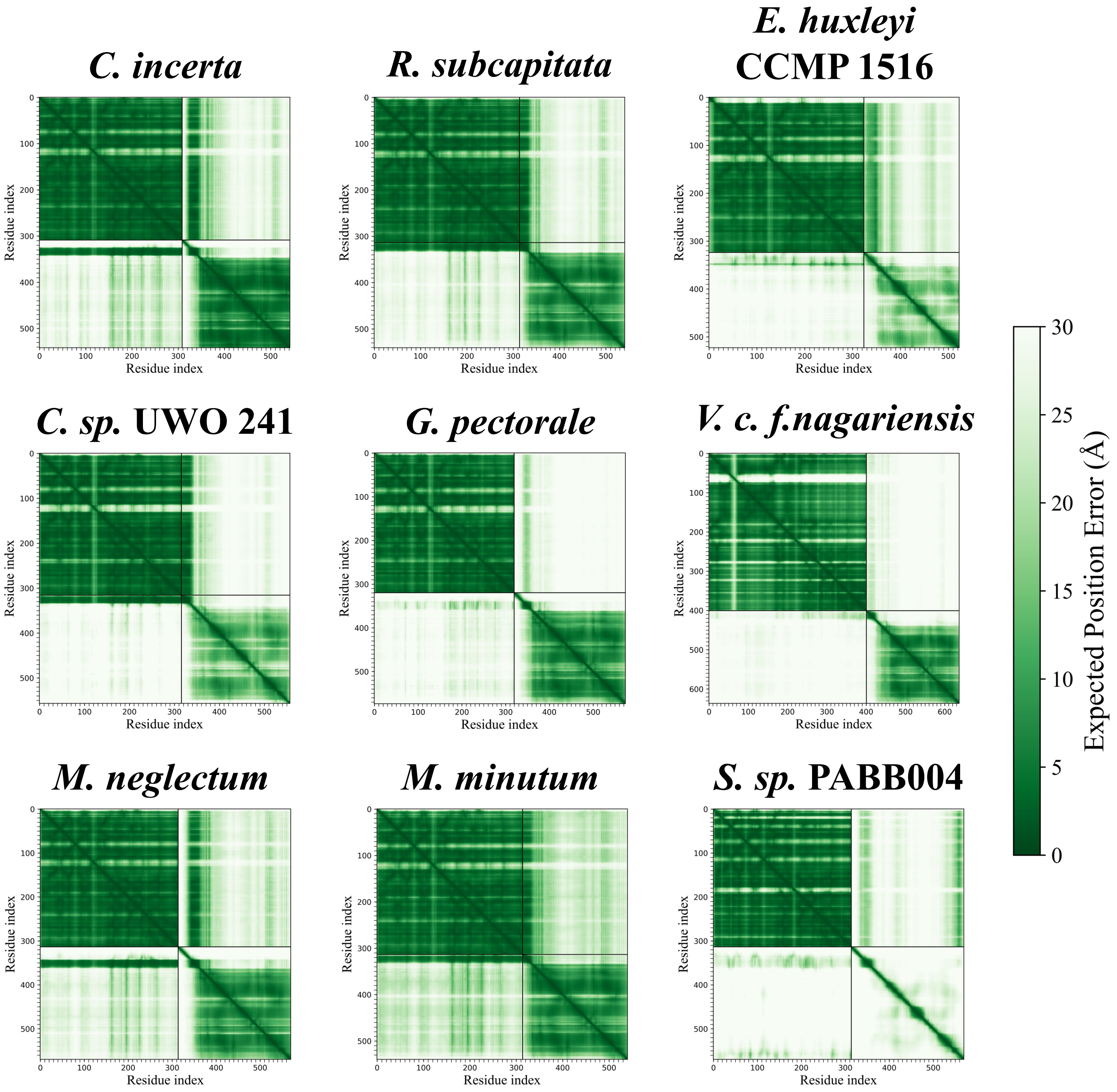


**Figure S14. PAE plots of FNR and Lhca proteins AlphaFold models.**

PAE heatmaps of the FNR-Lhca predicted complexes. Darker green indicates higher confidence in the relative positioning of residues. For each model, the two protein chains are concatenated and shown together.

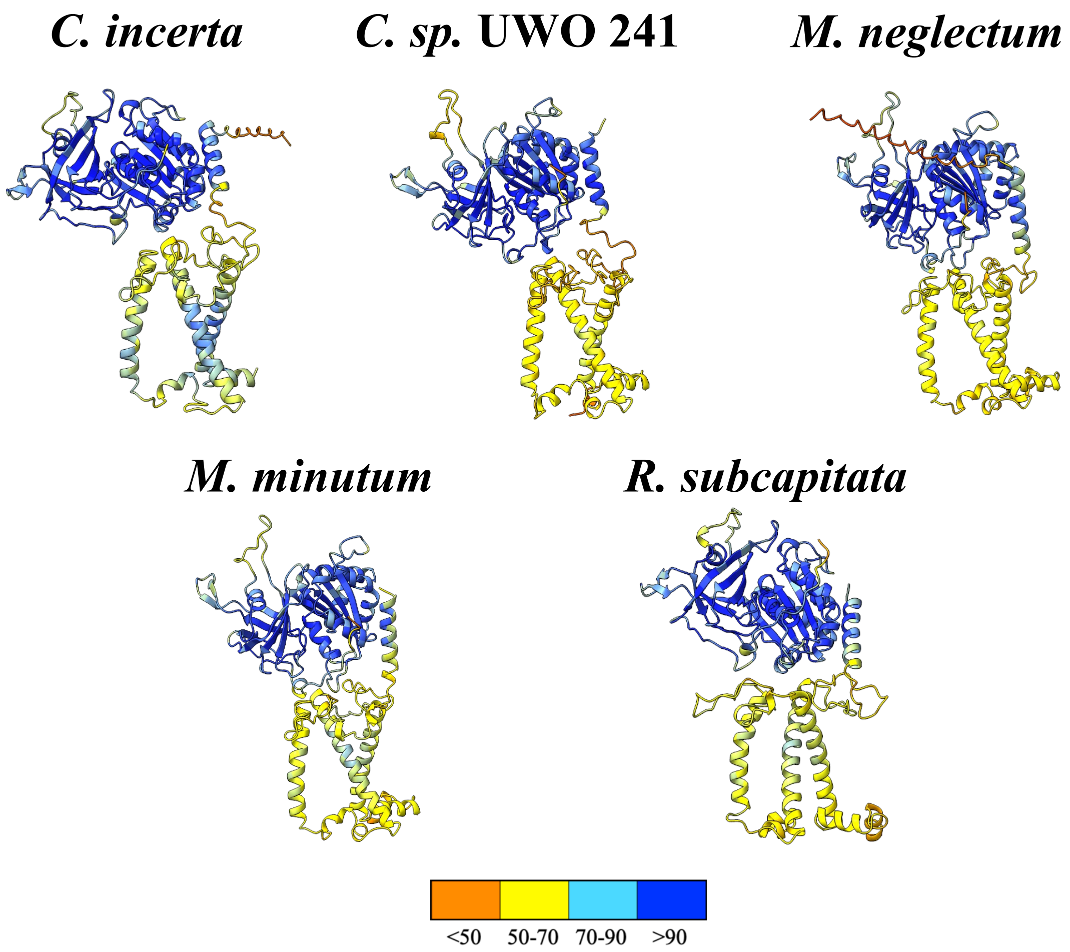


**Figure S15. FNR-Lhca proteins pairs modeled by AlphaFold and colored by pLDDT score.**

The models are of species whose Lhca proteins were identified as orthologs of *C. reinhardtii* Lhca4 (shown in **Fig. 5**) colored by per-residue pLDDT score, reflecting predicted local geometry accuracy: > 90 in blue, 70-90 in cyan, 50-70 in yellow and < 50 in orange.

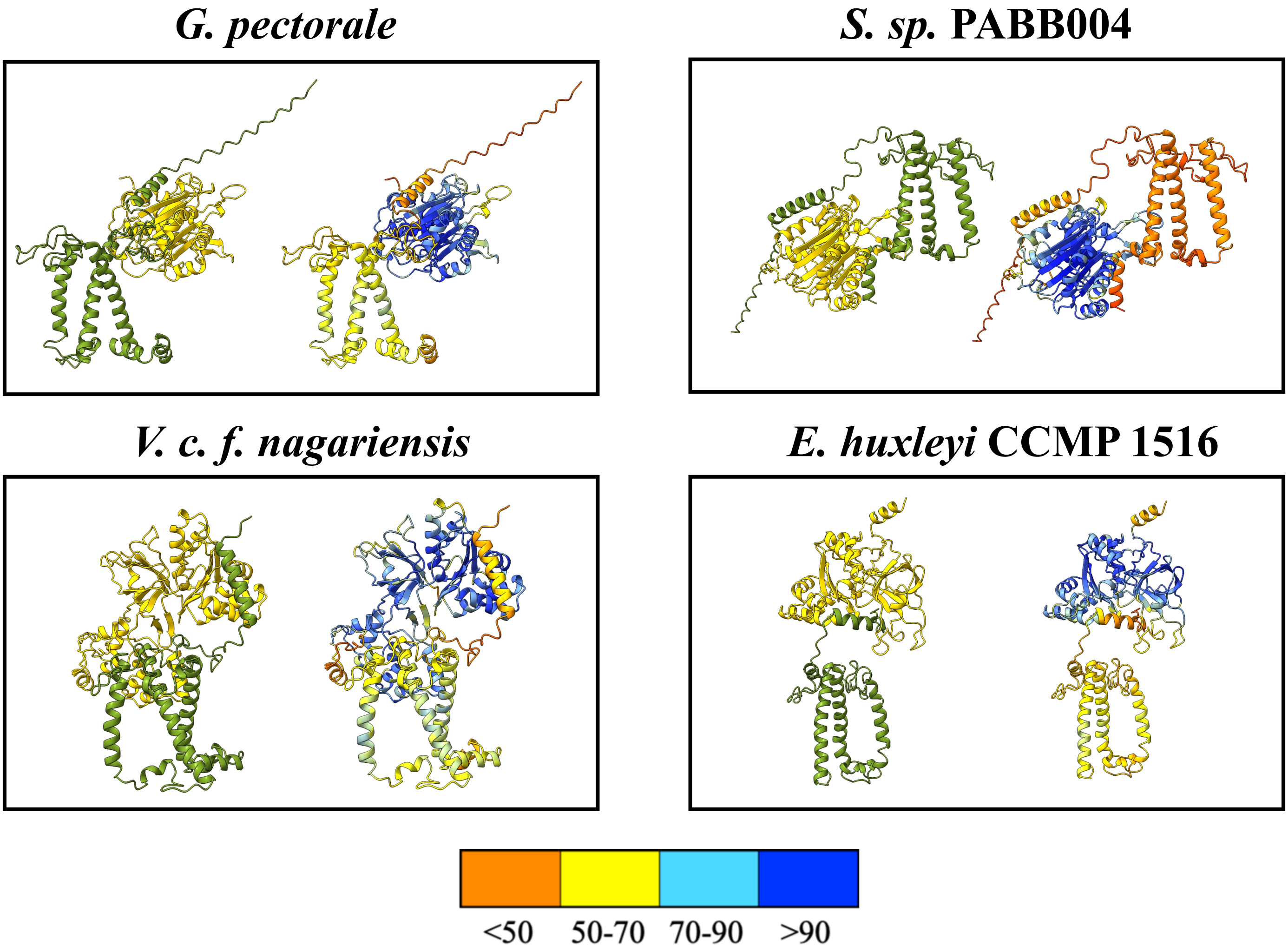


**Figure S16. Low-confidence AlphaFold models of Lhca-FNR complexes.**
Alphafold models of organisms identified in the search for C. reinhardtii Lhca4 homologs (green), in complex with their corresponding FNR proteins (gold). These are models that met all selection criteria, except for an adqequate ipTM score. The same model colored by pLDDT score is shown to its right: >90 (blue), 70-90 (cyan), 50-70 (yellow) and < 50 (orange), as indicated by the color bar.

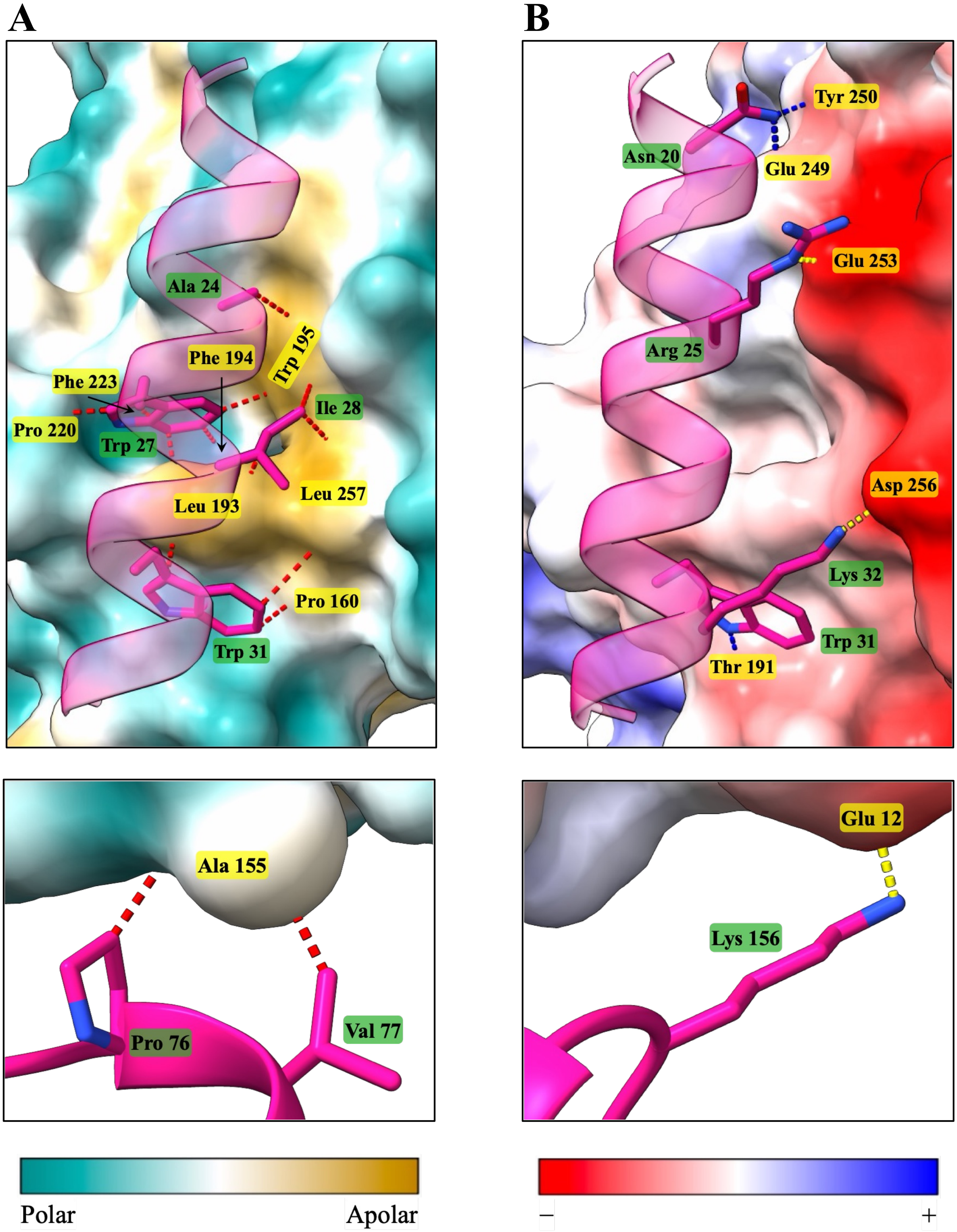


**Figure S17. Predicted FNR-Lhca interactions in *C. incerta*.**
Only the shortest distances between residue pairs are displayed, with FNR residues in yellow and Lhca residues in green.
**(A)** FNR surface colored by hydrophobicity, with cyan indicating hydrophilic (polar) regions and golden-brown indicating hydrophobic (apolar) regions. Hydrophobic contacts mediated by the N-terminal helix of Lhca (upper panel) are all <5 Å. Two additional hydrophobic interactions not involving the Lhca N-terminal helix are observed (lower panel). **(B)** FNR surface colored by electrostatic potential, ranging from red (negative) to blue (positive). Hydrogen bonds (blue distances) and salt bridges (yellow distances) mediated by the N-terminal helix of Lhca (upper panel) are all <3 Å. A single salt bridge not involving the Lhca N-terminal helix is observed at 2.7 Å (lower panel). No hydrogen bonds contributing to the FNR-Lhca interface are detected outside the N-terminal helix.

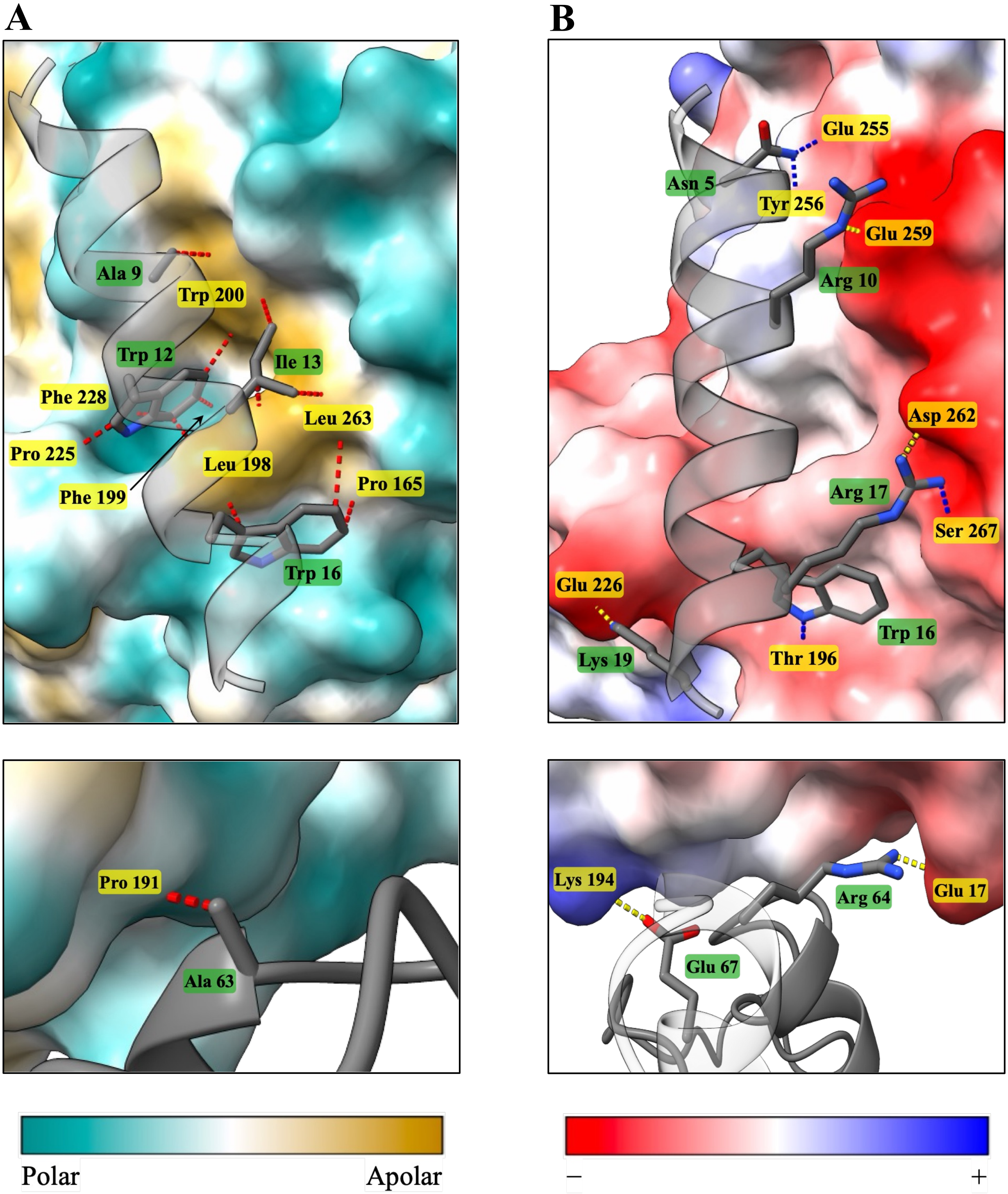


**Figure S18. Predicted FNR-Lhca interactions in *C. sp.* UWO 241.**
Only the shortest distances between residue pairs are displayed, with FNR residues in yellow and Lhca residues in green.
**(A)** FNR surfcae colored by hydrophobicity, with cyan indicating hydrophilic (polar) regions and golden-brown indicating hydrophobic (apolar) regions. Hydrophobic contacts mediated by the N-terminal helix of Lhca (upper panel) are all <5 Å. An additional hydrophobic interaction not involving the Lhca N-terminal helix is observed (lower panel) at 3.15 Å.
**(B)** FNR surface colored by electrostatic potential, ranging from red (negative) to blue (positive). Hydrogen bonds (blue distances) and salt bridges (yellow distances) mediated by the N-terminal helix of Lhca (upper panel) are all <3 Å A salt bridge not involving the Lhca N-terminal helix is observed at 2.784 Å (lower panel). No hydrogen bonds contributing to the FNR-Lhca interface are detected outside the N-terminal helix.

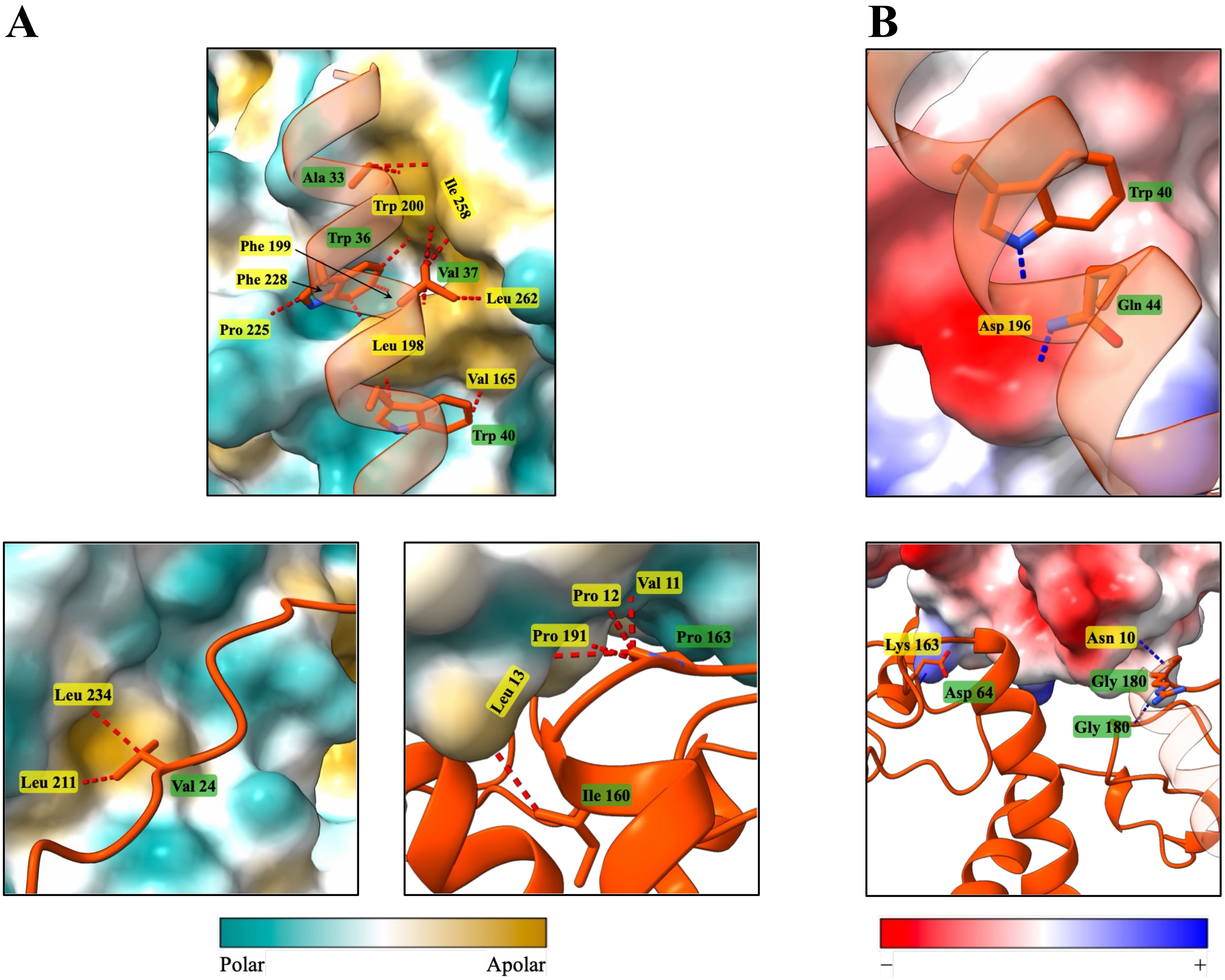


**Figure S19. Predicted FNR-Lhca interactions in *M. negelctum*.**

Only the shortest distances between residue pairs are displayed, with FNR residues in yellow and Lhca residues in green.

**(A)** FNR surfcae colored by hydrophobicity, with cyan indicating hydrophilic (polar) regions and golden-brown indicating hydrophobic (apolar) regions. Hydrophobic contacts mediated by the N-terminal helix of Lhca (upper panel) are all <5 Å. Additional hydrophobic contacts occur at an N-terminal segment prior to the helix (bottom-left panel) and subsequent to it (bottom-right panel).

**(B)** FNR surface colored by electrostatic potential, ranging from red (negative) to blue (positive). All are hydrogen bonds <3.5 Å occurring via the N-terminal helix of the Lhca protein (upper panel) or outside this region (lower panel).

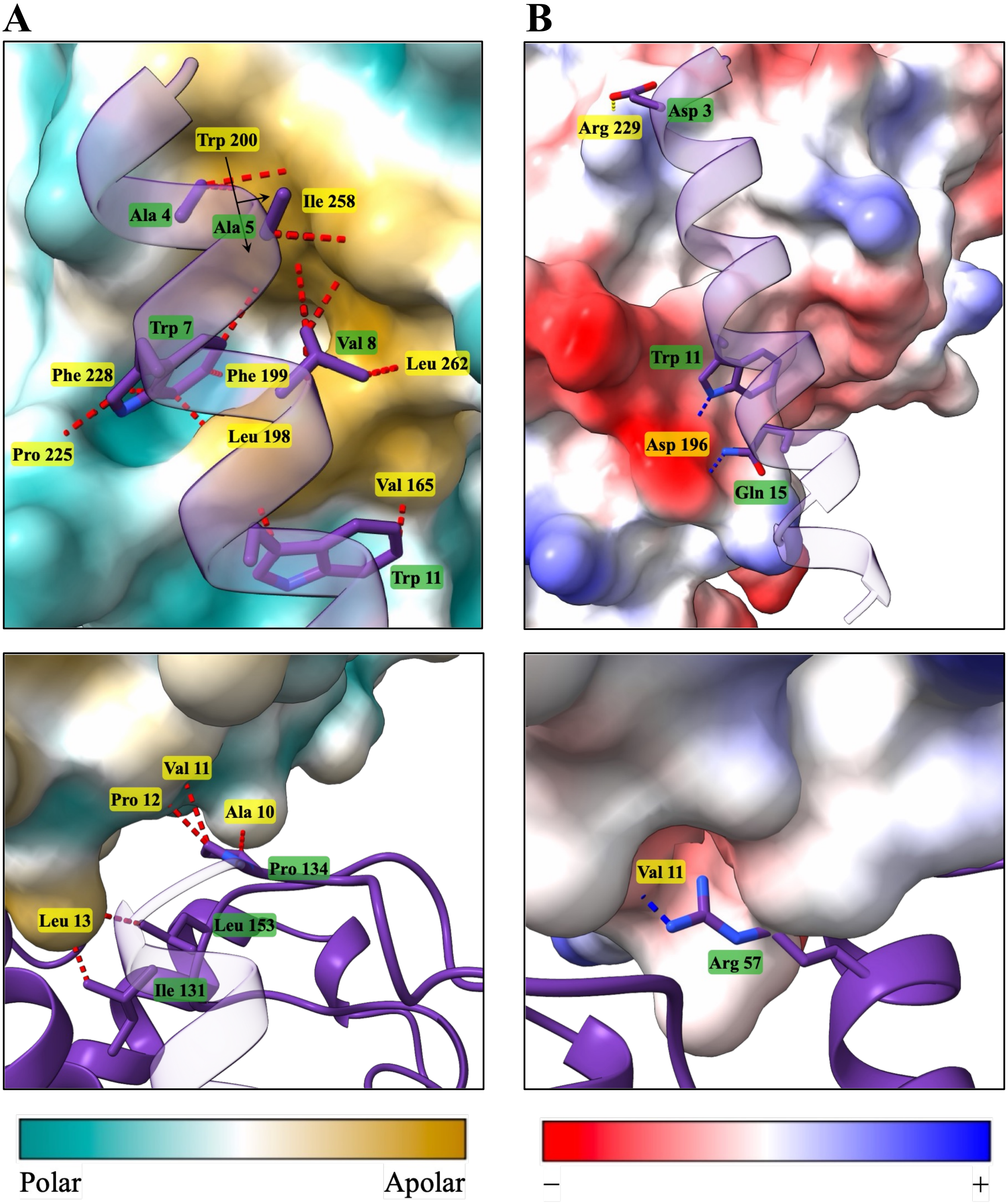


**Figure S20. Predicted FNR-Lhca interactions in *M. minutum.***

Only the shortest distances between residue pairs are displayed, with FNR residues in yellow and Lhca residues in green.

**(A)** FNR surfcae colored by hydrophobicity, with cyan indicating hydrophilic (polar) regions and golden-brown indicating hydrophobic (apolar) regions. Hydrophobic contacts mediated by the N-terminal helix of Lhca (upper panel) are all <5 Å. Additional hydrophobic interactions not involving the Lhca N-terminal helix are observed (lower panel), all <4.5 Å.

**(B)** FNR surface colored by electrostatic potential, ranging from red (negative) to blue (positive). Hydrogen bonds (blue distances) and salt bridges (yellow distances) mediated by the N-terminal helix of Lhca (upper panel) are all <3 Å. A single hydrogen bond not involving the Lhca N-terminal helix is observed at 2.8 Å (lower panel). No salt bridges contributing to the FNR-Lhca interface are detected outside the N-terminal helix.

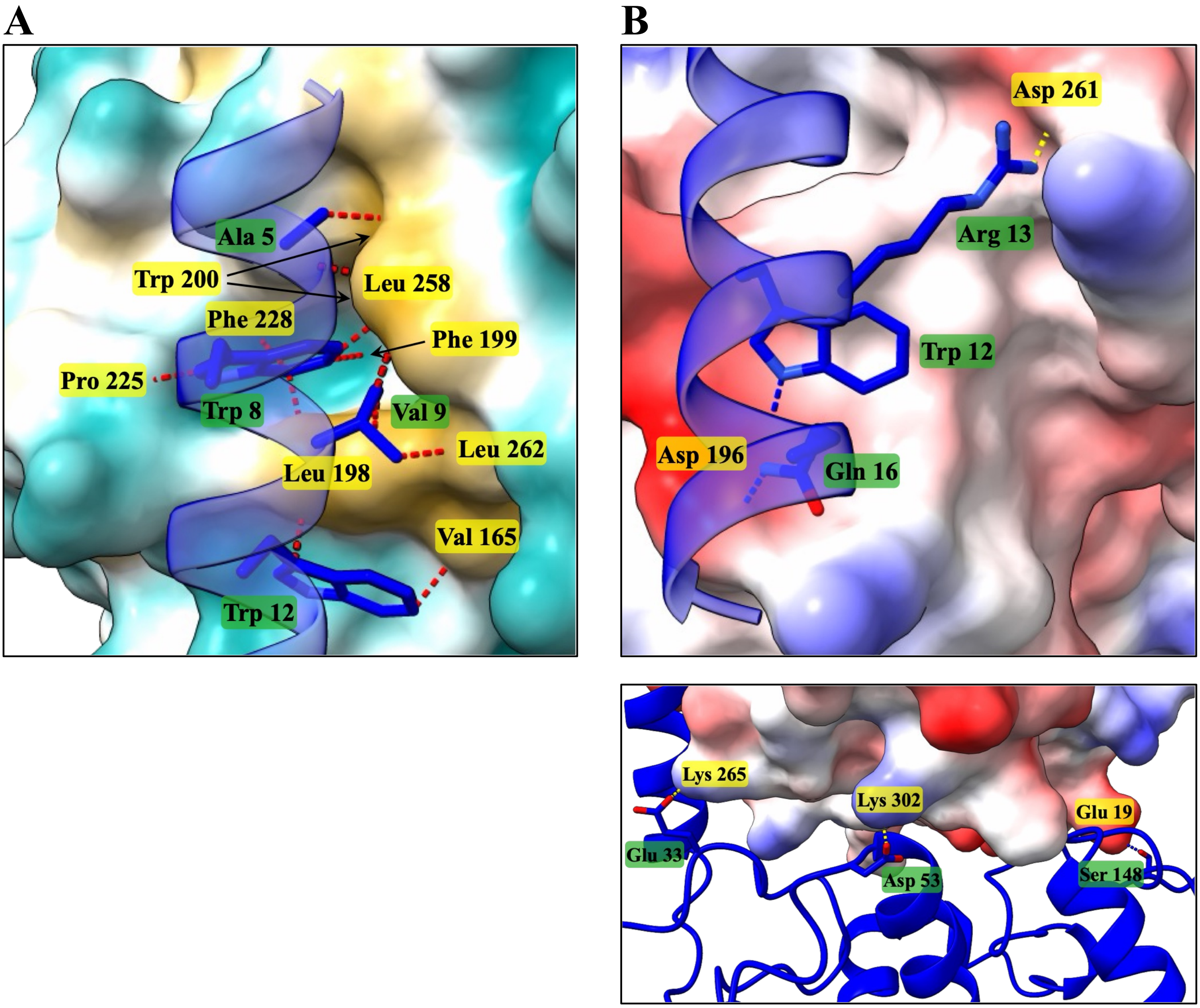


**Figure S21. Predicted FNR-Lhca interactions in *R. subcapitata*.**
Only the shortest distances between residue pairs are displayed, with FNR residues in yellow and Lhca residues in green.
**(A)** FNR surfcae colored by hydrophobicity, with cyan indicating hydrophilic (polar) regions and golden-brown indicating hydrophobic (apolar) regions. Hydrophobic contacts mediated by the N-terminal helix of Lhca (upper panel) are all <4.5 Å. There are no hydrophobic contacts outside this region.
**(B)** FNR surface colored by electrostatic potential, ranging from red (negative) to blue (positive). Hydrogen bonds (blue distances) and a salt bridge (yellow distances) mediated by the N-terminal helix of Lhca (upper panel) are all <3.5 Å. Additional hydrophobic contacts do not involve the N-terminal helix of the Lhca protein, all <3 Å (lower panel).

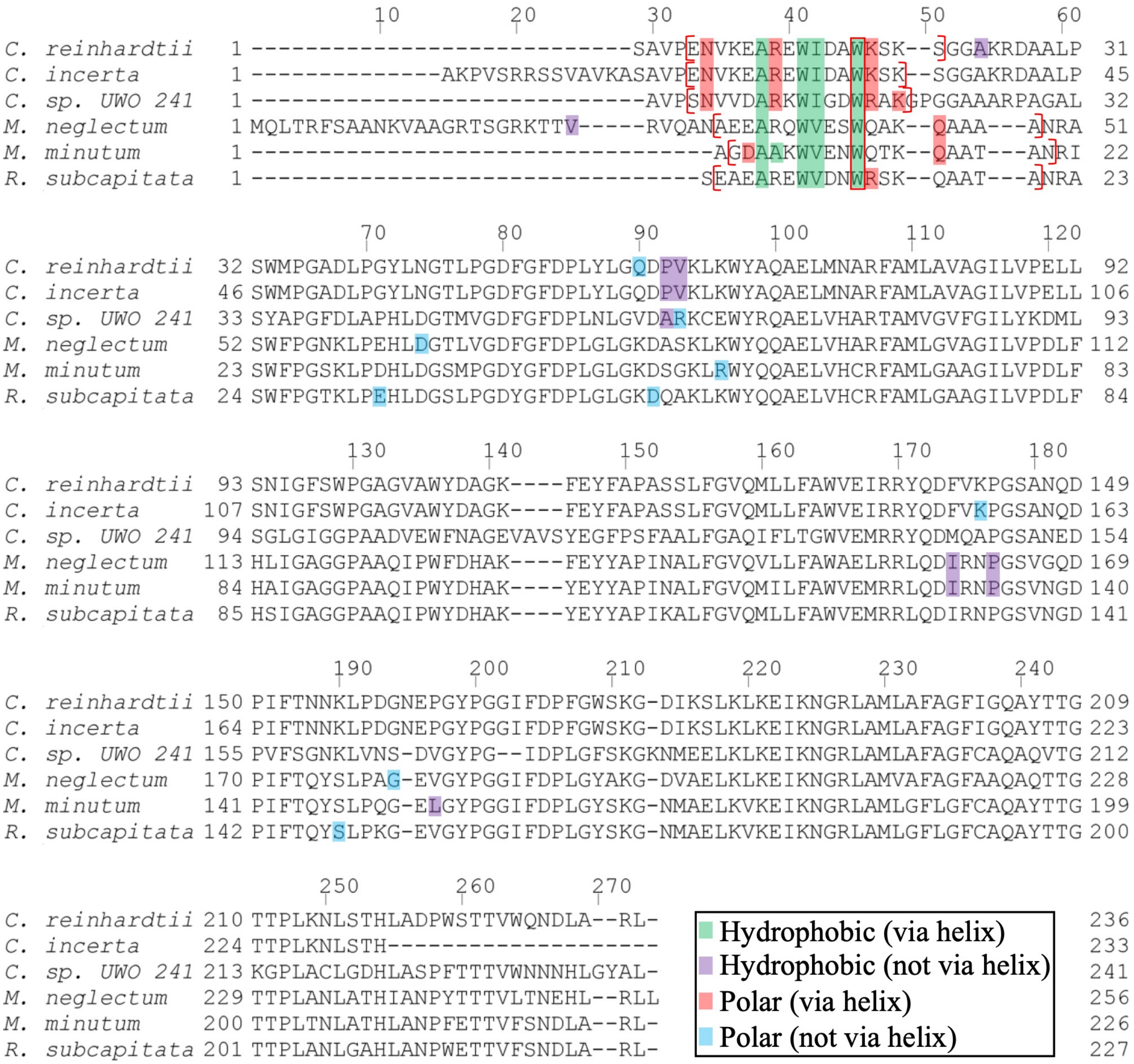


**Figure S22. Sequence alignment of Lhca4 homologs.**

Lhca protein sequences identified as homologs of *C. reinhardtii* Lhca4 were aligned together with it using MAFFT and visualized with JalView. Residues involved in hydrophobic interactions and van der Waals contacts are highlighted in green when located within the N-terminal helix and in purple when located outside it. Residues participating in polar interactions are highlighted in red within the N-terminal helix and in blue outside it. Residues involved in multiple types of interactions are framed according to their additional role. The N-terminal helix of each Lhca protein is indicated by red brackets.

```
                                10        20        30        40        50        60
C. reinhardtii   1 ATKASTAVTTDMSKRTVPTKLEEGEMPLNTYSNKAPFKAKVRSVEKITGPKATGETCHIII  61
C. incerta       1 -----------MSKRTVPTKLEEGEMPLNTYSNKAPFKAKIRSVETITGPKATGETCHIII  50
C. sp. UWO 241   1 ------ATAGKLAKASVPLKLEEGEMPLNTFSNKAPFVGRIKSVKRIVGPKATGETMHIII  55
M. neglectum     1 ------AATTTLTRTNVPLALEEGEMPLNTFSNKKPFTGTIRSVKRIVGPNATGETCDIVI  55
M. minutum       1 ------AAATAITRSAVPLKLEEGEMPLNTYSNKAPFKGTIKSVKRIVGPNATGETCDIVI  55
R. subcapitata   1 ------ATATSISRAAVPLKLEEGEMPLNTFSNKAPFKGTVKSVKRIVGPNATGETCDIVI  55

                         70        80        90       100       110       120
C. reinhardtii  62 ETEGKIPFWEGQSYGVIPPGTKINSKGKEVPHGTRLYSIASSRYGDDFDGQTASLCVRRAV 122
C. incerta      51 DTEGKIPFWEGQSYGVIPPGTKINSKGKEVPHGTRLYSIASSRYGDDFDGLTASLCVRRAV 111
C. sp. UWO 241  56 ETDGKIPFWEGQSYGVIPPGVKVNSRGKEVPHGVRLYSIASSRYGDTFDGMTTSLCVRRAN 116
M. neglectum    56 ETNGDIPYWEGQSYGVIPPGTKINSKGKEVPHGVRLYSIAASRYGDTFDGKTTTLCVRRAV 116
M. minutum      56 QTDGKIPFWEGQSYGVIPPGTKVNSKGKEVPYGVRLYSIASSRYGDEFDGMTTTLCVRRAV 116
R. subcapitata  56 ETNGKIPYWEGQSYGVIPPGTKVNSKGKEVPYGVRLYSIAASRYGDTFDGNTTTLCVRRAV 116

                       130       140       150       160       170       180
C. reinhardtii 123 YVDPETGKEDPAKKGLCSNFLCDATPGTEISMTGPTGKVLLLPADANAPLICVATGTGIAP 183
C. incerta     112 YVDPETGKEDPAKKGLCSNFLCDAKPGTEIMMTGPTGKVLLLPADTNAPLIMVATGTGIAP 172
C. sp. UWO 241 117 YWDEEMKADDPAKKGICSNFLCDAAPGTEIMMTGPAGKVLLLPDSPNTPVIMAATGTGIAP 177
M. neglectum   117 YVDPETGKEDPAKKGICSNFLCDAEPGTQIQMTGPAGKVLLLPESPKSVLICVATGTGIAP 177
M. minutum     117 FVDPETGKEDPAKKGICSNFLCDATPGTEISMTGPAGKVLLLPESPKSVLICVATGTGIAP 177
R. subcapitata 117 YVDPETGKEDPAKKGLCSNFLCDAAPGTEIMMTGPAGKILLMPESPKSVLICVATGTGIAP 177

                      190       200       210       220       230       240
C. reinhardtii 184 FRSFWRRCFIENVPSYKFTGLFWLFMGVANSDAKLYDEELQAIAKAYPGQFRLDYALSRE- 243
C. incerta     173 FRSFWRRCFIENVPSYKFTGLFWLFMGVANSDAKLYDEELQAIAKAYPSQFRLDYALSRE- 232
C. sp. UWO 241 178 FRSFWRRLFFENVPSYKYTGLFWLFMGAANSDGKLYDDELTQIAETYPENFRLDYALSREG 238
M. neglectum   178 FRSFWRRCFFENVPGWKFDGLFWLFMGVANSDSLLYEDEIQAIKATYPDNFRVDYALSRE- 237
M. minutum     178 FRSFWRRCFFESVPGWKFDGLFWLFMGVANSDSLLYDDEIKAISATAPDNFRVDYALSRE- 237
R. subcapitata 178 FRSFWRRCFYEDVPNWKFDGLFWLFMGVANSDSLLYEDEINAIKATYPDNFRVDYALSRE- 237

                     250       260       270       280       290       300
C. reinhardtii 244 QNNRKGGKMYIQDKVEEYADEIFDLLDN-GAHMYFCGLKGMMPGIQDMLERVAKEKGLNYE 303
C. incerta     233 QKNRKGGKMYIQDKVEEYADEIFDLLDN-GAHMYFCGLKGMMPGIQDMLERVAKEKGLNYE 292
C. sp. UWO 241 239 APNKRGGKMYIQDKMEEYADEIFDLLNNSGAHMYFCGLKGMMPGIQEMLERVCTEKGLKYE 299
M. neglectum   238 QKNKSGGKMYIQDKVEEYADIIFKLLDE-GAHIYFCGLKGMMPGITSMLERVAKAKGLNFE 297
M. minutum     238 QNNKSGGKMYIQDKVEEYADIIFKLLNE-GAHIYFCGLKGMMPGITSMLERVAKEKGINYE 297
R. subcapitata 238 QKNQSGGKMYIQDKVEEYSDLVFDLLNK-GAHIYFCGLKGMMPGIQSMLERVAKSKGLNFE 297

                    310       320
C. reinhardtii 304 EWVEGLKHKNQWHVEVY 320
C. incerta     293 EWVEGLKHKNQWHVEVY 309
C. sp. UWO 241 300 EFIEGLKHENRWHVEVY 316
M. neglectum   298 EFAEKLKHKNQWHVEVY 314
M. minutum     298 EFTEKLKHNNQWHVEVY 314
R. subcapitata 298 EYIEKLKHNNQWHVEVY 314
```

Hydrophobic (via helix)
Hydrophobic (not via helix)
Polar (via helix)
Polar (not via helix)

**Figure S23. Sequence alignment of orthogonal FNR proteins.**

FNR sequences from organisms harboring *C. reinhardtii* Lhca4 homologs were aligned with *C. reinhardtii* FNR using MAFFT and visualized in JalView. Residues mediating hydrophobic and van der Waals interactions with the Lhca N-terminal helix are highlighted in green, and those interacting elsewhere in purple. Residues involved in polar interactions with the N-terminal helix of Lhca proteins are highlighted in red, and those interacting with other Lhca regions in blue. Residues contributing to multiple interaction types are framed according to their additional role.

**A**

| | | | |
|---|---|---|---|
| *C. reinhardtii* | 1 | ENVKEAREWIDAWKSKS----- | 17 |
| *C. incerta* | 1 | ENVKEAREWIDAWKSK------ | 16 |
| *C. sp. UWO 241* | 1 | SNVVDARKWIGDWRAK------ | 16 |
| *M. neglectum* | 1 | A--EEARQWVESWQAKQAAAA- | 19 |
| *M. minutum* | 1 | ---GDAAKWVENWQTKQAATAN | 19 |
| *R. subcapitata* | 1 | E--AEAREWVDNWRSKQAATA- | 19 |
| *P. sativum Tic62* | 1 | ---LSPYTAYDDLKPPSSPSPT | 19 |
| *P. sativum Tic62* | 1 | ---LSPYAAYPDLKPPSSPSPS | 19 |
| *P. sativum Tic62* | 1 | ---LSPYAMYEDLKPPASPSPS | 19 |
| *A. thaliana Tic62* | 1 | ---LSPYASYEDLKPPTSPIPN | 19 |
| *O. sativa Tic62* | 1 | ---LSPYTRYEELKPPSSPTPS | 19 |
| *P. trichocarpa Tic62* | 1 | ---LSPYTAYEDLKPPTSPIPT | 19 |
| *A. thaliana Trol* | 1 | ---LSPYASYPDLKPPSSPMPS | 19 |
| *O. sativa subsp. japonica* | 1 | ---LSPYPNYPDLKPPSSPTPS | 19 |

**B**

| | | | |
|---|---|---|---|
| *C. reinhardtii* | 133 | EIRRYQDFVKPGSANQDPI | 151 |
| *C. incerta* | 147 | EIRRYQDFVKPGSANQDPI | 165 |
| *C. sp. UWO 241* | 138 | EMRRYQDMQAPGSANEDPV | 156 |
| *M. neglectum* | 153 | ELRRLQDIRNPGSVGQDPI | 171 |
| *M. minutum* | 124 | EMRRLQDIRNPGSVNGDPI | 142 |
| *R. subcapitata* | 125 | EMRRLQDIRNPGSVNGDPI | 143 |
| *P. sativum Tic62* | 5 | --TAYDDLKPPSSPSPT-- | 19 |
| *P. sativum Tic62* | 5 | --AAYPDLKPPSSPSPS-- | 19 |
| *P. sativum Tic62* | 5 | --AMYEDLKPPASPSPS-- | 19 |
| *A. thaliana Tic62* | 5 | --ASYEDLKPPTSPIPN-- | 19 |
| *O. sativa Tic62* | 5 | --TRYEELKPPSSPTPS-- | 19 |
| *P. trichocarpa Tic62* | 5 | --TAYEDLKPPTSPIPT-- | 19 |
| *A. thaliana Trol* | 5 | --ASYPDLKPPSSPMPS-- | 19 |
| *O. sativa subsp. japonica* | 5 | --PNYPDLKPPSSPTPS-- | 19 |

**Figure S24. Compaison of Lhca proteins to Tic62 and TROL.**

The FNR binding motifs of Tic62 and TROL from *Pisum sativum*, *Arabidopsis thaliana*, *Oryza sativa* and *Populus trichocarpa* (according to[23]) were aligned to **(A)** Lhca helix-forming residues and **(B)** full length algal Lhca proteins identified in this study. Residues in algal Lhca proteins predicted to interact with FNR and corresponding to the Tic62/TROL motifs are highlighted in red. In **(B)**, only the section aligning with the motifs and predicted to interact with FNR is shown.
