## Supplemental tables for "A Conserved Mechanism for Positioning Ferredoxin–NADP⁺ Reductase at Photosystem I in Green Algae"

| | PSI-LHCI-Fd (EMD-75304, PDB 10NJ) |
|---|---|
| **Data collection and processing** | |
| Magnification | 142,500 |
| Voltage (kV) | 300 |
| Electron exposure ($e^{-}/Å^{2}$) | 45 |
| Defocus range (μm) | 0.5 - 3 |
| Pixel size (Å) | 0.846 |
| Symmetry imposed | C1 |
| Initial particle images (no.) | 2,116,547 |
| Final particle images* (no.) | 169,538 |
| Map resolution (Å) | 2.20 |
| FSC threshold | 0.143 |
| **Refinement** | |
| Initial model used (PDB code) | 7ZQC |
| Model resolution (Å) | 2.21/2.52 |
| FSC threshold | 0.143,0.5 |
| Map sharpening *B* factor ($Å^{2}$) | 0 |
| Model composition | |
| Nonhydrogen atoms | 50,335 |
| Protein residues | 4246 |
| Ligands | 344 |
| Waters | 133 |
| *B* factors ($Å^{2}$) | |
| Protein | 71.34 |
| Ligand | 76.18 |
| Water | 68.92 |
| R.m.s. deviations | |
| Bond lengths (Å) | 0.005 |
| Bond angles (°) | 1.922 |
| **Validation** | |
| MolProbity score | 1.12 |
| Clashscore | 3.24 |
| Poor rotamers (%) | 0.24 |
| Ramachandran plot | |
| Favored (%) | 98.28 |
| Allowed (%) | 1.69 |
| Disallowed (%) | 0.02 |

**Table S1. Cryo EM data collection and refinement statistics.**
Summary of data acquisition, processing, reconstruction and validation parameters for each indicated dataset.

| Group | Subgroups | Tax ID |
|---|---|---|
| **Cyanobacteria** | - | 1117 |
| **SAR (Stramenopiles, Alveolata, and Rhizaria)** | - | 2698737 |
| **Hacrobia** | Haptophytes | 2830 |
| | Telonemids (Telonemida) | 589438 |
| | Centrohelids | 193537 |
| | Biliphytes (Picobiliphytes) | 419944 |
| | Katablepharids (Katablepharida) | 339960 |
| | Cryptomonads | 3027 |
| **Unikonts** | Opisthokonts | 33154 |
| | Amoebozoa | 554915 |
| | Apusomonads | 172820 |
| | Breviates (Breviatea) | 1401294 |
| **Excavata** | Kinetoplastids | 5653 |
| | Diplonemids (Diplonemida) | 191814 |
| | Euglenids | 3035 |
| | Heterolobosea | 5752 |
| | Jakobids (Jakobida) | 556282 |
| | Oxymonads | 66288 |
| | Parabasalids | 5719 |
| | Retortamonads | 193075 |
| | Diplomonads | 5738 |
| | Malawimonads (Malawimonadidae) | 136087 |
| **Chlorophyta** | - | 3041 |
| **Glaucophyta** | - | 38254 |
| **Rhodophyta** | - | 2763 |
| **Embryophyta** | - | 3193 |
| **Charophyta** | - | 3146 |

**Table S2. Lineages screened in the search for Lhca4 homologs.** Taxonomic identifiers (Tax ID) of lineages and their respective subgroups included in the BLASTP search for *C. reinhardtii* Lhca4 homologs.

| Organism | Tax ID |
|---|---|
| *Chlamydomonas incerta* | 51695 |
| *Chlamydomonas sp. UWO 241* | 1653778 |
| *Monoraphidium neglectum* | 145388 |
| *Monoraphidium minutum* | 39955 |
| *Raphidocelis subcapitata* | 307507 |
| *Gonium pectorale* | 33097 |
| *Volvox carteri f. nagariensis* | 3068 |
| *Scenedesmus sp.* PABB004 | 2742167 |
| *Emiliania huxleyi* CCMP1516 | 280463 |

**Table S3. Taxonomic identifiers of organisms used to retrieve FNR sequences.**
The taxonomic identifiers (Tax ID) of organisms whose N-termini are homologous to *C. reinhardtii* Lhca4 and are predicted by AlphaFold to form an alpha-helix are listed here. They were utilized in BLASTP searches aimed to find their FNR sequences, by using *C. reinhardtii* FNR sequence as the query.

| Organism | Taxonomic group | Mean pLDDT | Mean pLDDT of helix | ipTM | pTM |
|---|---|---|---|---|---|
| *Chlamydomonas reinhardtii* | Green algae | 86.85 | 92.23 | 0.87 | 0.63 |
| *Chlamydomonas incerta* | Green algae | 88.86 | 95.03 | 0.9 | 0.64 |
| *Chlamydomonas sp.* UWO 241 | Green algae | 82.06 | 96.7 | 0.9 | 0.62 |
| *Monoraphidium neglectum* | Green algae | 82.12 | 90.68 | 0.88 | 0.64 |
| *Monoraphidium minutum* | Green algae | 85.92 | 84.39 | 0.85 | 0.67 |
| *Raphidocelis subcapitata* | Green algae | 86.56 | 92.37 | 0.89 | 0.64 |
| *Gonium pectorale* | Green algae | 80.07 | 47.15 | 0.2 | 0.57 |
| *Volvox carteri f. nagariensis* | Green algae | 82.05 | 57.12 | 0.18 | 0.58 |
| *Scenedesmus sp.* PABB004 | Green algae | 66.05 | 52.52 | 0.23 | 0.58 |
| *Emiliania huxleyi* CCMP1516 | Coccolithophores | 79.25 | 48.71 | 0.53 | 0.64 |

**Table S4. AlphaFold models pTM and ipTM scores.**
Shown are the pTM and iPTM scores of the AlphaFold models generated in this work, signifying the confidence of the predictions. Green colored organisms' names are ones with ipTM scores of above 0.8, their models are shown in **Fig. 5**. The rest have ipTM score of below 0.6, their models are shown in **Fig. S16**. The PAE plots are shown in **Fig. S14**.
